# Broadening a SARS-CoV-1 neutralizing antibody for potent SARS-CoV-2 neutralization through directed evolution

**DOI:** 10.1101/2021.05.29.443900

**Authors:** Fangzhu Zhao, Meng Yuan, Celina Keating, Namir Shaabani, Oliver Limbo, Collin Joyce, Jordan Woehl, Shawn Barman, Alison Burns, Xueyong Zhu, Michael Ricciardi, Linghang Peng, Jessica Smith, Deli Huang, Bryan Briney, Devin Sok, David Nemazee, John R. Teijaro, Ian A. Wilson, Dennis R. Burton, Joseph G. Jardine

## Abstract

The emergence of SARS-CoV-2 underscores the need for strategies to rapidly develop neutralizing monoclonal antibodies that can function as prophylactic and therapeutic agents and to help guide vaccine design. Here, we demonstrate that engineering approaches can be used to refocus an existing neutralizing antibody to a related but resistant virus. Using a rapid affinity maturation strategy, we engineered CR3022, a SARS-CoV-1 neutralizing antibody, to bind SARS-CoV-2 receptor binding domain with >1000-fold improved affinity. The engineered CR3022 neutralized SARS-CoV-2 and provided prophylactic protection from viral challenge in a small animal model of SARS-CoV-2 infection. Deep sequencing throughout the engineering process paired with crystallographic analysis of an enhanced antibody elucidated the molecular mechanisms by which engineered CR3022 can accommodate sequence differences in the epitope between SARS-CoV-1 and SARS-CoV-2. The workflow described provides a blueprint for rapid broadening of neutralization of an antibody from one virus to closely related but resistant viruses.

## INTRODUCTION

Following the emergence of severe acute respiratory syndrome coronavirus 2 (SARS-CoV-2), a massive effort was initiated to repurpose existing or discover new antibodies against SARS-CoV-2 to use as research tools, diagnostics, and as direct medical countermeasures for prophylactic and therapeutic indications. Early repurposing efforts screened monoclonal antibodies that had previously been isolated from 2003 SARS-CoV-1 (sometimes designated as SARS-CoV) and 2008 MERS-CoV survivors against SARS-CoV-2 (Tian et al., 2020), but none of the existing antibodies were able to efficiently neutralize this novel virus. In parallel, multiple groups worked to isolate antibodies from SARS-CoV-2 infected humans or from animals immunized with SARS-CoV-2 spike (S) proteins (Hansen et al., 2020; Ju et al., 2020; Pinto et al., 2020; Robbiani et al., 2020; Rogers et al., 2020; Wrapp et al., 2020). Although various approaches were used in the antibody discovery process, most utilized an antigen-specific B cell sorting strategy to isolate binding antibodies followed by an *in vitro* neutralization assay to identify the neutralizing subset. This antibody discovery process has been widely used to identify HIV neutralizing antibodies and extensively refined and streamlined over the last decade to the point where it is possible to progress from biological samples to recombinantly produced antibodies ready for validation in 1-2 weeks (Huang et al., 2013, 2014; Rogers et al., 2020; Sok et al., 2014). The two major bottlenecks in this process currently are: 1) access to high quality peripheral blood mononuclear cell (PBMC) samples for antigen-specific B cell sorting and 2) the identification of lead therapeutic candidates that have both the desired neutralization function and biochemical developability properties that are amenable to large-scale manufacturing and formulation. Access to PBMCs was particularly onerous in the very early stages of the COVID-19 outbreak when donor samples were not available in the United States and Europe because of shipping and/or biosafety restrictions.

Here we explore a hybrid “refocusing” approach that is a blend between conventional discovery and repurposing, where an existing neutralizing antibody is engineered to target a related, but resistant virus. As a case study, we selected CR3022, a SARS-CoV-1 neutralizing monoclonal antibody isolated in 2006 from a convalescent donor (ter Meulen et al., 2006). At the onset of the pandemic, CR3022 received considerable attention because it was shown to cross-react with SARS-CoV-2 (Tian et al., 2020). Several groups tested whether CR3022 could neutralize SARS-CoV-2, and most observed either no neutralization or only partial neutralization at the highest antibody concentration (Anand et al., 2020; Atyeo et al., 2021; Manenti et al., 2020; Wu et al., 2020b; Yuan et al., 2020a; Zhou et al., 2020), although one group did report neutralizing activity in their assay (Huo et al., 2020). A crystal structure revealed that CR3022 recognizes an epitope outside of the ACE2 binding site that is highly conserved between SARS-CoV-1 and SARS-CoV-2 (Yuan et al., 2020a) with only four amino acid differences located in or around the CR3022 epitope. Reversion of one of these four mutations, P384A, was shown to be responsible for the 100-fold reduction in binding affinity for CR3022 to the novel SARS-CoV-2 (Wu et al., 2020a). These findings are consistent with other documented examples of viral escape from neutralizing antibodies (nAbs), where small changes in the antibody epitope are sufficient to reduce antibody binding below the requisite threshold to achieve effective neutralization (Bates et al., 2014). To address this, we wanted to explore whether engineering approaches can be used to retarget the SARS-CoV-1 nAb CR3022 to the corresponding epitope on SARS-CoV-2.

## RESULTS

### Engineering of SARS-CoV-1 nAb CR3022 to efficiently recognize SARS-CoV-2

To engineer CR3022 variants with higher affinity for SARS-CoV-2 S protein, we utilized a rapid antibody affinity maturation strategy that we developed called **<u>S</u>**ynthetic **<u>A</u>**ntibody **<u>M</u>**aturation by multiple **<u>P</u>**oint **<u>L</u>**oop library **<u>E</u>**n**<u>R</u>**ichments (SAMPLER). Libraries are generated around a starting antibody sequence by introducing single mutations within the complementarity-determining region (CDR) loops. These variant libraries are then displayed on the surface of yeast as molecular Fab and screened with fluorescence-activated cell sorting (FACS) to isolate clones with improved affinity for the target antigen (Figure 1A). During the library generation, the diversity is introduced into the CDR loops using predefined oligo pools (Li et al., 2018), where each mutation is explicitly synthesized so the library is free of unwanted mutations like cysteines or methionines and mutations that would introduce N-linked glycan motifs. Each CDR loop library contains 100-200 unique variants, depending on the length of the loop, and when the CDR1/2/3 libraries are combined, the resulting combinatorial library contains 3-4 million unique sequences, each with up to 3 mutations from the parental CR3022 heavy chain (HC) or light chain (LC) sequence (Figure 1A). This strategy efficiently samples a large theoretical search space around the starting antibody sequence and allows for the screening of synergistic mutations between CDR loops, as it has been shown that affinity enhancing mutations can be destabilizing and require compensatory stabilizing mutations (Julian et al., 2017). Overall, this rational CDR synthesis reduces the theoretical diversity by 6-fold compared to using NNK by removing the redundancy of the overrepresented amino acids and removing unwanted mutations (Table S1).

**Figure 1.**
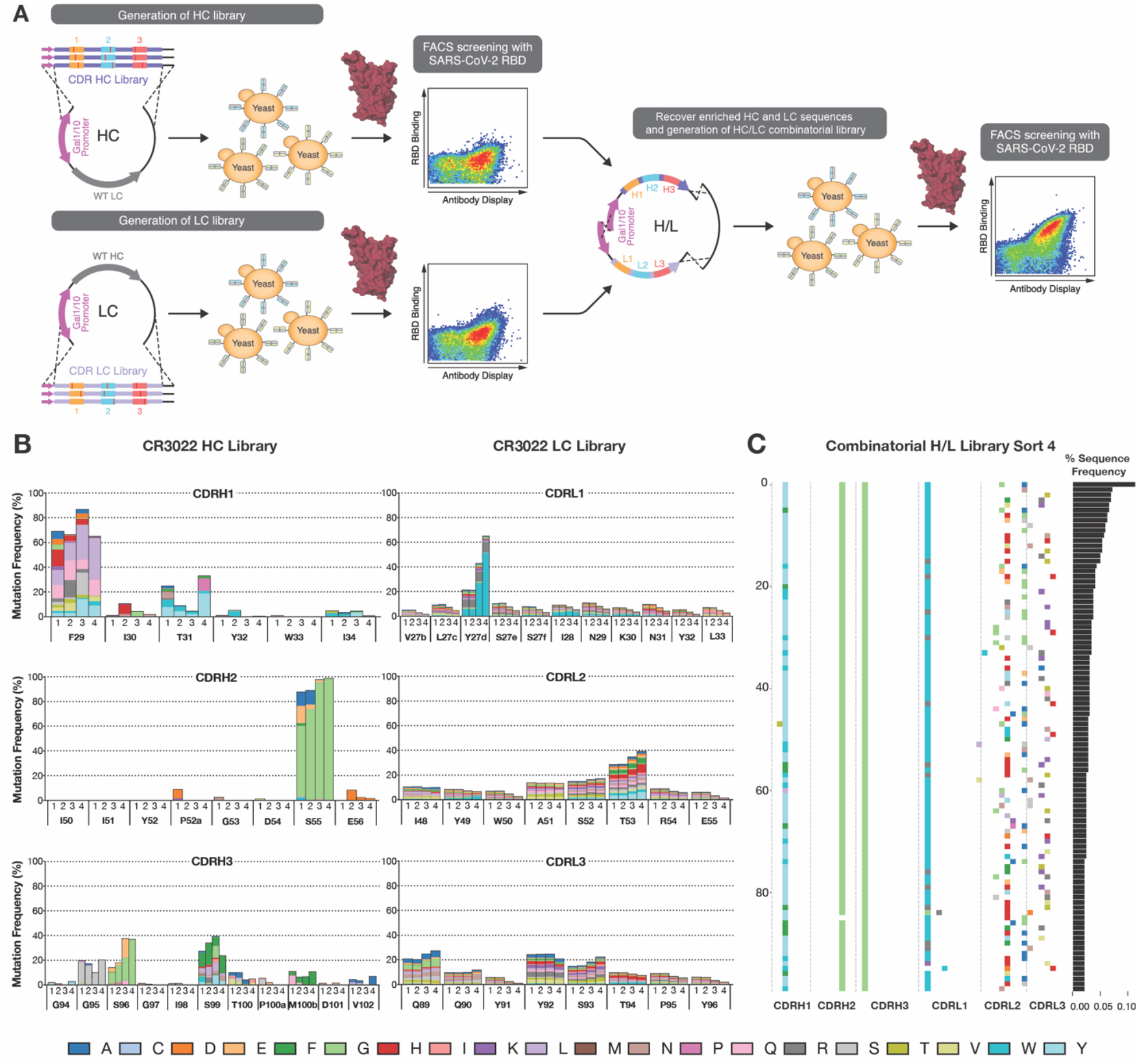
Engineering CR3022 to increase binding affinity to SARS-CoV-2 RBD using SAMPLER. (**A**) A synthetic CR3022 antibody library with single mutation at each CDR loop was displayed as molecular Fab on the surface of yeast cells. The CR3022 HC library with up to three mutations was paired with the original LC while LC library was paired with the original HC. After FACS selection by SARS-CoV-2 RBD, the HC library and LC library were amplified and combined into combinatorial H/L library and further selected for high binding clones by SARS-CoV-2 RBD. Enriched clones with high binding affinities were reformatted and expressed as human IgG. (**B**) After each round of selection with a total of four FACS sorts, plasmid DNA from each sort was prepped and deep sequenced. Enriched mutations for each residue at CDR loop in HC library (left) and LC library (right) relative to parental CR3022 sequence were analyzed and colored according to the key. (**C**) Locations of mutations (colored according to the key) seen in the top 100 most frequent sequences recovered from long-read next-generation sequencing of enriched H/L pairs in the CR3022 combinatorial H/L library after sort 4.

Initially, two libraries were generated—one library with mutations in the HC paired with unmodified LC (HC library) and one library with mutations in the LC that was paired with the unmodified HC (LC library) and displayed on the surface of yeast (Figure 1A). Each library was sorted four times against SARS-CoV-2 receptor binding domain (RBD) to enrich for variants with higher affinity for the protein (Figure S1). In sorts 1,2 and 4, cells were labeled with non-saturating concentrations of biotinylated SARS-CoV-2 RBD and the top 5-10% of RBD-binding cells, normalized for Fab surface display, were collected to enrich for HC or LC sequences with increased affinity for SARS-CoV-2 RBD (Figure S2). In sort 3, a negative selection was used to deplete polyreactive clones, where cells were labeled with a biotinylated preparation of detergent solubilized HEK293 membrane proteins (Figure S1) (Xu et al., 2013). After the four selections, the antibody display vectors from the HC and LC libraries were harvested and the region encoding the heavy and light chain was amplified and combined into a new combinatorial heavy and light chain library (H/L library) that sampled diversity in both chains (Figure 1A). The H/L library was screened with the same four-round selection protocol to identify the optimal combination of mutations in the heavy and light chains. In total, it took just under one month to complete all three rounds of SAMPLER optimization (Figure S1).

Following each round of SARS-CoV-2 selection, plasmid DNA encoding the HC and LC regions was amplified and deep sequenced. The HC library showed a strong preference for an S55G mutation (Kabat numbering) in CDRH2, appearing in 57% of sequences after sort 1 and 98% of sequences after sort 4 (Figure 1B). In CDRH3, an S96G mutation was observed in 36% of sequences. The mutations in CDRH1 were largely localized to the F29 and T31 positions, with 65% and 33% enriched mutation frequency respectively, but multiple mutations were allowed at each one of those positions. The LC library showed strong enrichment for a tyrosine to tryptophan mutation at position Y27_d_ residue in CDRL1, enriching to 52% of the total reads after sort 4 (Figure 1B). CDRL2 and CDRL3 showed no strong enrichments across any of the selections.

In the combinatorial H/L library, three mutations were further enriched in CR3022 HC: 98% T31W in CDRH1, 99% of S55G in CDRH2, and 100% S96G in CDRH3 (Figure S3). Nevertheless, the LC remained more diverse in the H/L library, with the Y to W mutation further enriched to 71% of the population at the Y27_d_ position (Figure S3). In addition to deep sequencing of the individual chains, PacBio sequencing was used to evaluate the recovered heavy/light pairs from sort 4. The overall frequencies of mutations closely matched what was observed in the individual chains, with a high degree of convergence on the heavy chain and a significant level of diversity still present in the light chain, with no clear evidence of evolutionary coupling within the selected H/L chain pairings (Figure 1C).

### Binding and neutralizing activity of eCR3022 for SARS-CoV-2

From the sequences recovered after the final H/L library sort, we selected 25 engineered CR3022 antibodies, named engineered (e) eCR3022.1 through eCR3022.25, to reformat for expression as human IgG1 for in-depth characterization. Because of the nature of how the H/L library was constructed and the fact that there was more observed convergence in the HC, the selected variants utilized only 5 unique HCs, while all 25 LCs were unique (Table S2). The affinities for SARS-CoV-1 RBD of all eCR3022 Abs remained the same or slightly improved compared to the parental CR3022, despite not having been included in the optimization process (Figures 2A, 2B; Table S2). The binding affinities of all eCR3022 antibodies against monomeric SARS-CoV-2 RBD increased between 100 to 1000-fold compared to the parental CR3022 by surface plasmon resonance (SPR), with equilibrium dissociation constant (K_D_) values ranging from 16 pM to 312 pM (Figures 2A, 2C; Table S2). The on-rate of the antibodies was nearly the same as the parental CR3022, with the affinity increases coming through a reduction in the antibody off-rate (Figure 2C, Table S2). The ELISA binding activities of the eCR3022 variants against SARS-CoV-2 S and RBD were also improved (Figure S4), showing that this increased binding to SARS-CoV-2 RBD also translated to the functional S protein. All engineered antibodies showed negligible polyreactive binding across our nonspecific antigen panels (Figure S5A, S5B) and were monodispersed with analytical SEC column retention times similar to other clinical mAbs (Figure S5C), indicating the engineering had minimal impact on the favorable biochemical properties of the parental CR3022.

**Figure 2.**
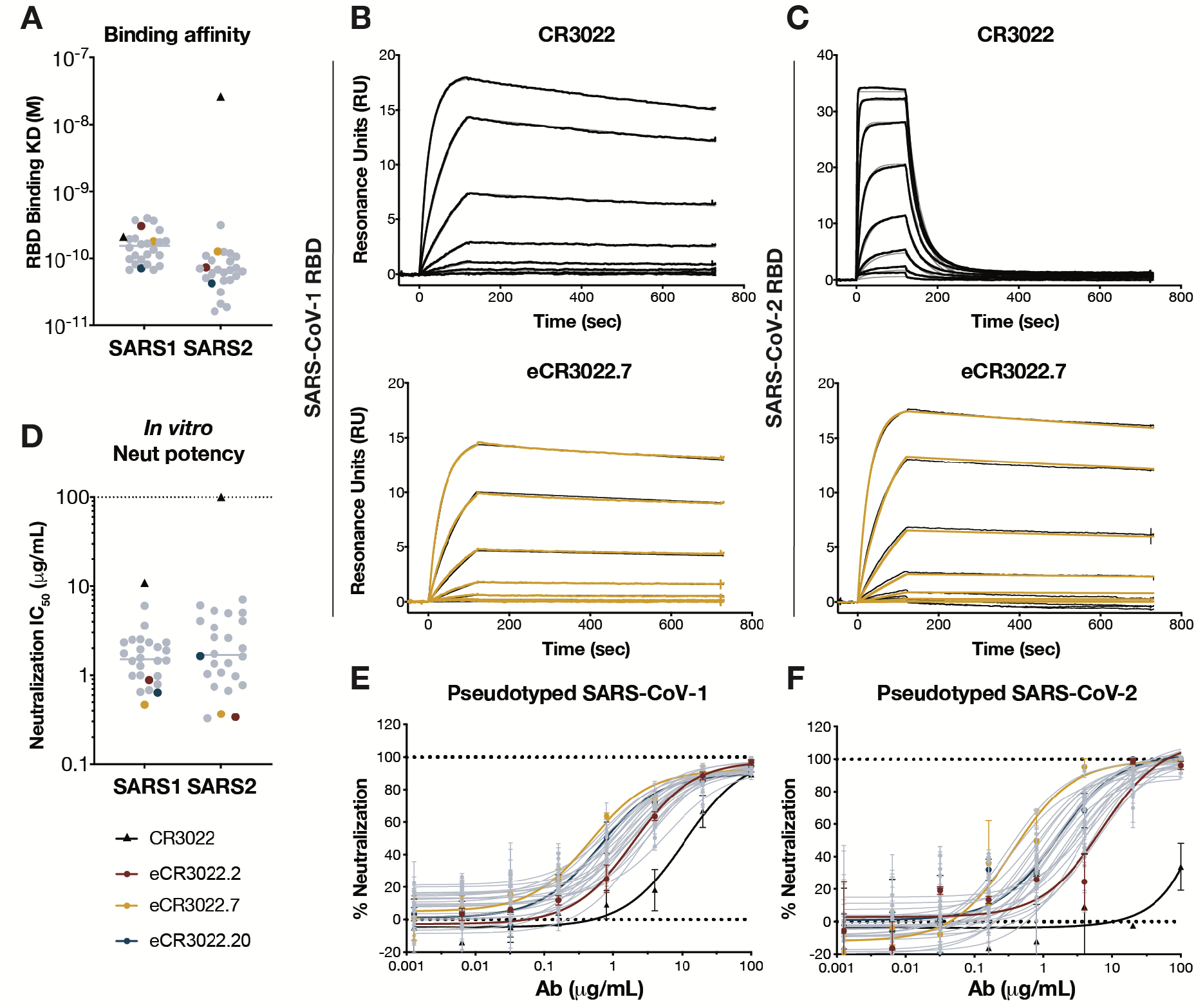
Engineered eCR3022 variants with over 100-fold improved binding affinity for SARS-CoV-2 RBD potently neutralize SARS-CoV-2. **(A)** Binding affinity (KD) of parental and enhanced eCR3022 antibodies against SARS-CoV-1 and SARS-CoV-2 RBD by surface plasmon resonance (SPR). Parental CR3022 is colored in black and represented as a triangle, eCR3022 variants are in grey whereas eCR3022.2, eCR3022.7, eCR3022.20 are highlighted in colors according to the key. (**B**) SPR curves of parental CR3022 (top) and eCR3022.7 (bottom) binding to SARS-CoV-1 RBD. (**C**) SPR curves of parental CR3022 (top) and eCR3022.7 (bottom) binding to SARS-CoV-2 RBD. Antibodies were captured via Fc-capture to an anti-human IgG Fc antibody and varying concentrations of SARS-CoV-1 or SARS-CoV-2 RBD were injected using a multi-cycle method. Association and dissociation rate constants calculated through a 1:1 Langmuir binding model using the BIAevaluation software. (**D**) Neutralization IC_50_s of parental CR3022 and eCR3022 antibodies against SARS-CoV-1 and SARS-CoV-2 pseudoviruses. (**E-F**) Neutralization curves of parental CR3022 and eCR3022 antibodies against SARS-CoV-1 (**E**) and SARS-CoV-2 (**F**).

To evaluate the effect of the affinity gains on antibody neutralization, CR3022 and all 25 eCR3022 variants were tested using an MLV-based pseudovirus system (Rogers et al., 2020).

All variants showed enhanced neutralization of SARS-CoV-1, with the most potent neutralizing at 0.5 μg/mL compared to the parental CR3022 that neutralizes at 10.9 μg/mL (Figures 2D, 2E). In our assay, CR3022 failed to neutralize SARS-CoV-2 at a maximum antibody concentration of 100 μg/mL (Figures 2D, 2F). In contrast, all eCR3022 antibodies were able to neutralize pseudotyped SARS-CoV-2 with a median IC_50_ of 1.6 μg/mL, with the most potent neutralizing SARS-CoV-2 exhibiting an IC_50_ of 0.3 μg/mL (Figures 2D, 2F). We further tested the neutralization activity of eCR3022.7, eCR3022.10 and eCR3022.20 against the emerging variants of concern B.1.1.7 (with N501Y mutation on RBD) and B.1.351 (with K417N, E484K, N501Y mutations on RBD) variants (Tegally et al.), where the mutations are located in the receptor binding site on the RBD and are outside the CR3022 epitope. As expected, eCR3022 antibodies neutralized both viral variants with similar IC_50_ to the wildtype virus SARS-CoV-2 (Figure S6). Lastly, the eCR3022 variants were able to neutralize authentic SARS-CoV-2 virus while CR3022 failed to neutralize, confirming the neutralization against authentic virus (Figure S7).

### Structural analysis of eCR3022.20 complexed to SARS-CoV-2 RBD

To understand the molecular features of the affinity-matured eCR3022 antibodies that confer potent neutralization against SARS-CoV-2, we determined a crystal structure of eCR3022.20 in complex with SARS-CoV-2 RBD and Fab CC12.3 (Yuan et al., 2020b) (to aid in crystallization) to 2.85 Å resolution and compared its binding with CR3022 (Yuan et al., 2020a) (Figure 3A; Figure S8; Table S3). Two copies of the eCR3022.20-RBD-CC12.3 complex were found in the crystal asymmetric unit. eCR3022.20 binds SARS-CoV-2 RBD via the same epitope as CR3022 through a nearly identical angle of approach (Yuan et al., 2020a) (Figure 3A). Furthermore, the RBD conformation bound by both antibodies are almost identical, except for two regions of the RBD (residues 365-370 and 384-390) that are displaced in eCR3022.20 compared to CR3022 (Figure 3B). However, residues in both of these regions (RBD-Y369, F377, and P384) form a hydrophobic pocket that interact with CDRH1 of these antibodies (Figures 3C-3F). V_H_ T31 in CDRH1 of CR3022 (Figure 3C) is substituted by a bulky hydrophobic, aromatic residue W31, which stacks with RBD residues Y369, F377, and P384 and strengthens the interaction with this hydrophobic pocket in the RBD (Figures 3D, 3E). The T31W substitution induces a 2.4-Å shift in the RBD around residues 365-370, as well as a 1.4-Å shift of residues 384-390 (Figure 3F). Previously, we showed that residue 384 is an important epitope residue for CR3022 binding, where a P384A mutation conferred an approximate 100-fold affinity improvement for CR3022 (Wu et al., 2020a). Here, we show that a substitute paratope residue targeting this area of the epitope is able to contribute to an improvement in antibody binding and further highlights the importance of P384 as a key epitope residue for CR3022.

**Figure 3.**
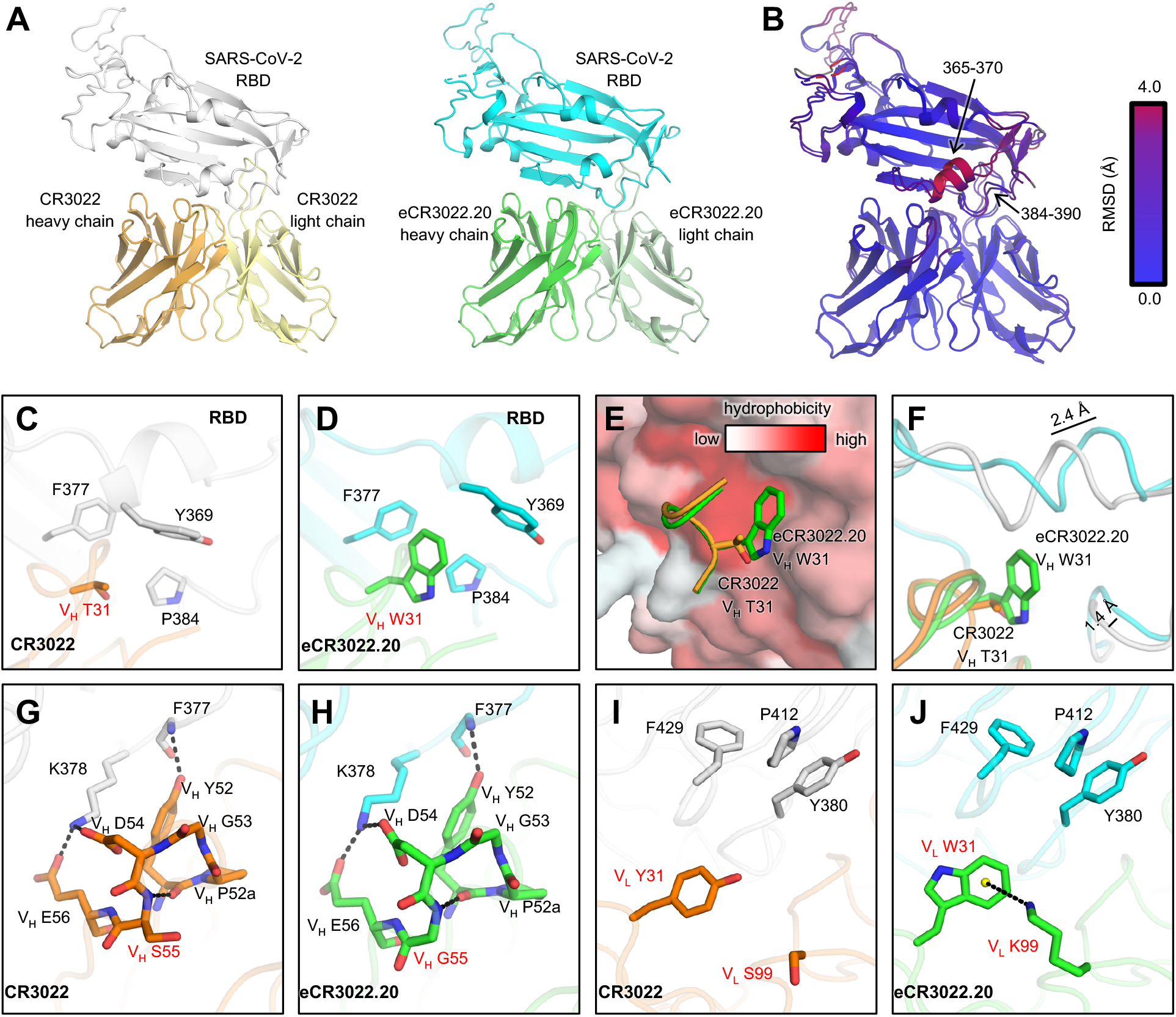
Crystal structure of eCR3022.20 in complex with SARS-CoV-2 RBD. Our previous crystal structure of SARS-CoV-2 RBD in complex with CR3022 is shown for comparison (PDB: 6W41) (Yuan et al., 2020a). Hydrogen bonds and salt bridges are represented by black dashed lines. Residues that differ between the two antibodies are highlighted in red letters. (**A**) CR3022 and eCR3022.20 bind to SARS-CoV-2 RBD via the same binding mode. Left, crystal structure of SARS-CoV-2 RBD (white) in complex with CR3022 (heavy and light chains are shown in orange and light yellow, respectively). Right, crystal structure of SARS-CoV-2 RBD (cyan) in complex with eCR3022.20 (heavy and light chains are shown in green and olive, respectively). Antibody constant domains are omitted here for clarity. Antibody CC12.3 (Yuan et al., 2020b) that was used to aid in the crystallization of the RBD/eCR3022.20 complex is shown in Figure S8. (**B**) Superimposition of structures RBD/CR3022 and RBD/eCR3022.20. Structural differences are color-coded by their Root Mean Square Deviation (RMSD). (**C-F**) Comparison between the paratope V_H_ T31 in CR3022 and its counterpart V_H_ W31 in eCR3022.20. (**E**) V_H_ T31/W31 interact with a hydrophobic pocket in the SARS-CoV-2 RBD. (**G-H**) Interactions between CDR H2 of (**G**) CR3022 and (**H**) eCR3022.20 with the SARS-CoV-2 RBD. (**I-J**) V_L_ Y31/S99 in CR3022 are substituted by W31/K99 in eCR3022.20. (**J**) V_L_ W31 and K99 in eCR3022.20 form a cation-π interaction. The 6-carbon aromatic ring center is represented by a yellow sphere.

In addition to CDRH1, CDRH2 of both CR3022 and eCR3022.20 interact with SARS-CoV-2 RBD (Figures 3G, 3H). For both antibodies, V_H_ Y52 forms a hydrogen bond with the backbone amide of RBD-F377. V_H_ D54 and E56 clamp onto RBD-K378 with two salt bridges (Figures 3G, 3H). CDRH2 in both antibodies form a type-IV β (Lewis et al., 1973) at the CDR tip, where V_H_ S55 (i+3) in CR3022 is substituted by a glycine in most sorted antibody variants during the process of affinity maturation (Figure 1B; Figure S9). Glycine is frequently found in β turns due to the lack of a side chain that allows higher conformational flexibility. Indeed, glycine is the most frequent residue at the i+3 position of a type-IV β turn, suggesting that it is energetically favored at this position as it can also more readily adopt a positive phi value as found here compared to S55 (Guruprasad and Rajkumar, 2000). For the light-chain residues, V_L_ Y31 and S99 are mutated to W31 and K99 in eCR3022.20 (Figures 3I, 3J). While V_L_ W31 retains hydrophobic interactions with RBD-Y380, P412, and F429, V_L_ K99 is able to form a cation-π interaction with W31, which stabilizes the interaction between CDRs L1 and L3 and may reduce the entropy of its interaction with the RBD.

### Protection by eCR3022 against SARS-CoV-2 challenge in a small animal model

Upon developing a CR3022 variant capable of neutralizing SARS-CoV-2, we sought to evaluate whether this engineered antibody could provide prophylactic protection against viral challenge in the Syrian hamster model. Based on the neutralization and biophysical data, eCR3022.7 was selected as our initial candidate antibody to evaluate. Groups of six hamsters received an intraperitoneal infusion 10 mg, 2 mg, 0.5 mg, or 0.125 mg of eCR3022.7 per animal to evaluate dose-dependent protection. Another group received 10 mg of the parental CR3022 antibody and a control group received 10 mg of an anti-dengue isotype matched antibody. Three days post infusion, sera were collected from each animal to determine antibody titer at the time of viral challenge. Animals were then challenged with 1 x 10^5^ plaque forming units (PFU) of SARS-CoV-2 (USA-WA1/2020) by intranasal administration. The hamsters were weighed daily as a measure of disease due to infection and sera were collected from all animals at the conclusion of the experiment on day 7. Hamsters have been shown to clear SARS-CoV-2 infection after 7 days (Sia et al., 2020), so a replicate of the original experiment was performed with six additional animals in which lung tissue was collected on day 4 to accurately measure replicative viral load amongst the groups (Figure 4A).

**Figure 4.**
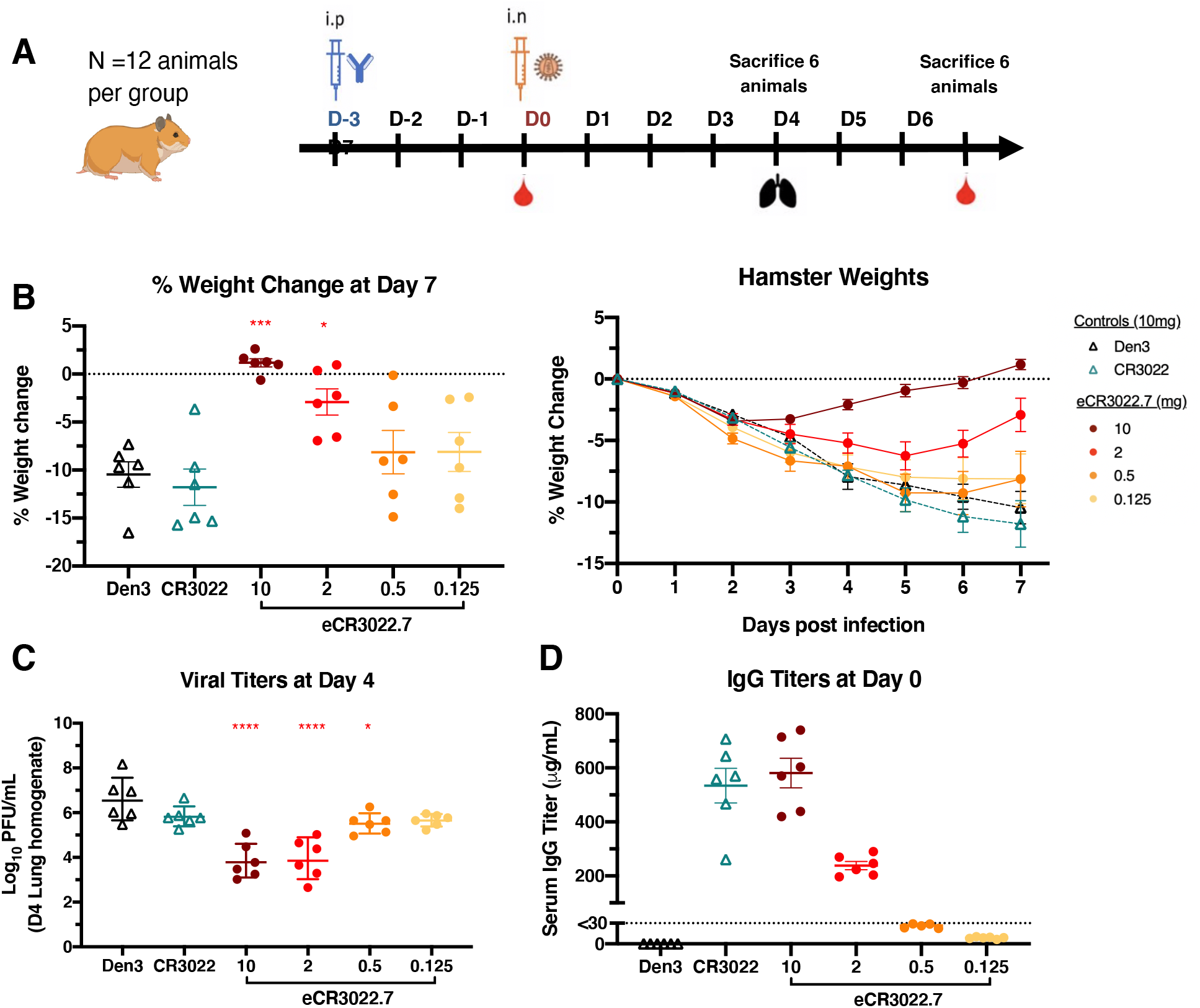
A non-neutralizing mAb can be engineered to become a nAb against SARS-CoV-2 and protects against weight loss and lung viral replication in Syrian hamsters. (**A**) engineered mAb eCR3022.7 was administered at a starting dose of either 10 mg per animal, 2 mg per animal, 0.5 mg per animal or 0.125 mg per animal. Control animals received 10 mg of Den3 or 10 mg of original mAb CR3022. Each group of 12 animals was challenged intranasally (i.n.) 72 hours after infusion with 1 × 10^5^ PFU of SARS-CoV-2. Serum was collected at the time of challenge (day 0) and upon completion of the experiment (day 7). Animal weight was monitored as an indicator of disease progression. Six hamsters were sacrificed on day 4 and lung tissue was collected for viral burden assessment and the remaining hamsters were sacrificed on day 7. (**B**) Percent weight change was calculated from the day of infection (day 0) for all animals. (**C**) Viral load, as quantitated by live virus plaque assay on Vero E6 cells from lung tissue homogenate. Error bars represent geometric standard deviations of the geometric mean. (**D**) Serum titers of the passively administered mAb, as assessed by ELISA at the time of challenge (72 hours after intraperitoneal (i.p.) administration). Statistical significance (p < 0.05) of groups in comparison to Den3 IgG control group were calculated by Ordinary One-Way ANOVA test using Graph Pad Prism 8.0. For weights and serum titers, error bars represent group average with standard error of the mean.

The groups receiving the highest doses of eCR3022.7 of 10 mg or 2 mg were largely protected from viral challenge, exhibiting either no weight loss or 3% weight loss, respectively, at the end of the 7-day experiment (Figure 4B; Figure S10). In contrast, the group that received 10 mg of the original CR3022 antibody lost 11% of body weight, comparable to the control group that received 10 mg of Den3. The groups that received 0.5 mg or 0.125 mg eCR3022.7 on average lost slightly less weight than the control group but the difference was not significant (Table S4). Lung viral titers were assessed from the second group of animals by plaque assay (Figure 4C). Equivalent viral loads were measured between the control Den3 and original CR3022 groups with average of 4.0 x 10^6^ PFU/mL and 6.8 x 10^5^ PFU/mL, respectively (Table S4). In contrast, the groups that received 10 mg or 2 mg doses of eCR3022.7 had 7.0 x 10^4^ PFU/mL and 8.7 x 10^4^ PFU/mL, respectively, more than 2 logs lower than the Den3 control. The reduced SARS-CoV-2 viral lung titers and maintenance of body weight following challenge in the animals receiving either 10 mg or 2 mg doses prophylactically of eCR3022.7 demonstrate that the *in vitro* neutralization potency translates to *in vivo* protection. Serum antibody concentrations were measured both at the time of challenge and the end of the study. The 10 mg eCR0322.7 and 10 mg CR3022 groups had an equivalent amount of antibody present at the time of injection as well as equivalent decay by day 7, confirming that the observed protection differences are not attributable to different pharmacokinetic properties of the two antibodies (Figure 4D, Figure S10B).

## DISCUSSION

Antibody refocusing serves as a novel approach to generate neutralizing antibodies against novel viruses or viral variants, provided that a neutralizing antibody against a closely related virus is available. In this case study, we successfully re-engineered a SARS-CoV-1 nAb, CR3022, so that it now potently neutralized both SARS-CoV-1 and SARS-CoV-2. eCR3022 variants with >1000-fold enhanced affinity for SARS-CoV-2 RBD were generated while maintaining their biochemically favorable developability profile. The enhanced affinity conferred the ability to potently neutralize SARS-CoV-2 and protect from viral challenge in a small animal model. In fact, the eCR3022 variants now neutralize SARS-CoV-2 more potently than the parental CR3022 neutralizes SARS-CoV-1. These findings are consistent with previous studies showing a relationship between antibody/antigen binding affinity for S protein and neutralization potency. However, this is the first instance that we are aware of where an antibody has been specifically retargeted or optimized to a novel related virus. Although this refocusing strategy likely will not be effective when the antibody epitope is substantially different between viruses (Figure S11), the potential pool of starting antibodies continues to expand as more neutralizing antibodies are discovered against different viruses. We also note that the use of on-chip DNA synthesis to rationally produce the CDR libraries used in our SAMPLER optimization allowed us to efficiently cover a large portion of the theoretical search space within the antibody paratope, effectively achieving these affinity gains in a two-step process. Given that the affinity engineering described can be done in less than one month and does not require access to PBMC samples from immunized or infected donors or structural information on the antibody/antigen interaction, the approach could be used in conjunction with conventional antibody discovery as a rapid response to future outbreaks of pandemic concern.

## ACKNOWLEDGEMENTS

We thank Laurence Cagnon at The Scripps Research Institute for coordinating with BSL-3 experiments. We thank Scripps Genomic Core facility for the help of deep sequencing. We thank Raiees Andrabi at The Scripps Research Institute for providing SARS-CoV-1 S protein. This work was partially funded by IAVI with the generous support of USAID and other donors; a full list of IAVI donors is available at www.iavi.org. This work was supported by the Bill and Melinda Gates Foundation OPP1170236 and INV-004923 INV (I.A.W., D.R.B.). D.H. and D.N. were supported by R01AI132317. Use of the SSRL, SLAC National Accelerator Laboratory, is supported by the U.S. Department of Energy, Office of Science, Office of Basic Energy Sciences under Contract No. DE-AC02-76SF00515. The SSRL Structural Molecular Biology Program is supported by the DOE Office of Biological and Environmental Research, and by the National Institutes of Health, National Institute of General Medical Sciences (including P41GM103393).

## AUTHOR CONTRIBUTIONS

F.Z., C.K., J.G.J. designed the experiments. J.G.J. designed the synthetic antibody library, F.Z., S.B. and M.J.R. performed the yeast library display and FACS selection. F.Z. and S.B. prepped the yeast library DNA and deep sequenced. J.S. analyzed the deep sequencing results and C.J. analyzed the PacBio sequencing results. F.Z., O.L., S.B., and A.B. expressed and purified the antibodies. J.W. performed the SPR assay. F.Z., O.L. S.B., and A.B. carried out the pseudovirus neutralization assays. S.P. and D.H. generated SARS-CoV-2 mutant virus constructs. N.S and D.H. performed authentic SARS-CoV-2 neutralization assay. M.Y. and O.L. expressed and purified the recombinant SARS-CoV-2 S and RBD proteins. F.Z., S.B. A.B. performed binding assays, and biophysical analysis assays. M.Y., X.Z., and I.A.W. crystallized eCR3022.20 and performed structure determination and analysis. C.K. and N.S. performed the hamster protection study and measured the viral load. F.Z., M.Y., C.K., I.A.W., D.R.B. and J.G.J. wrote the manuscript, and all authors reviewed and edited the manuscript.

## SOM Tables

**Table S1.**
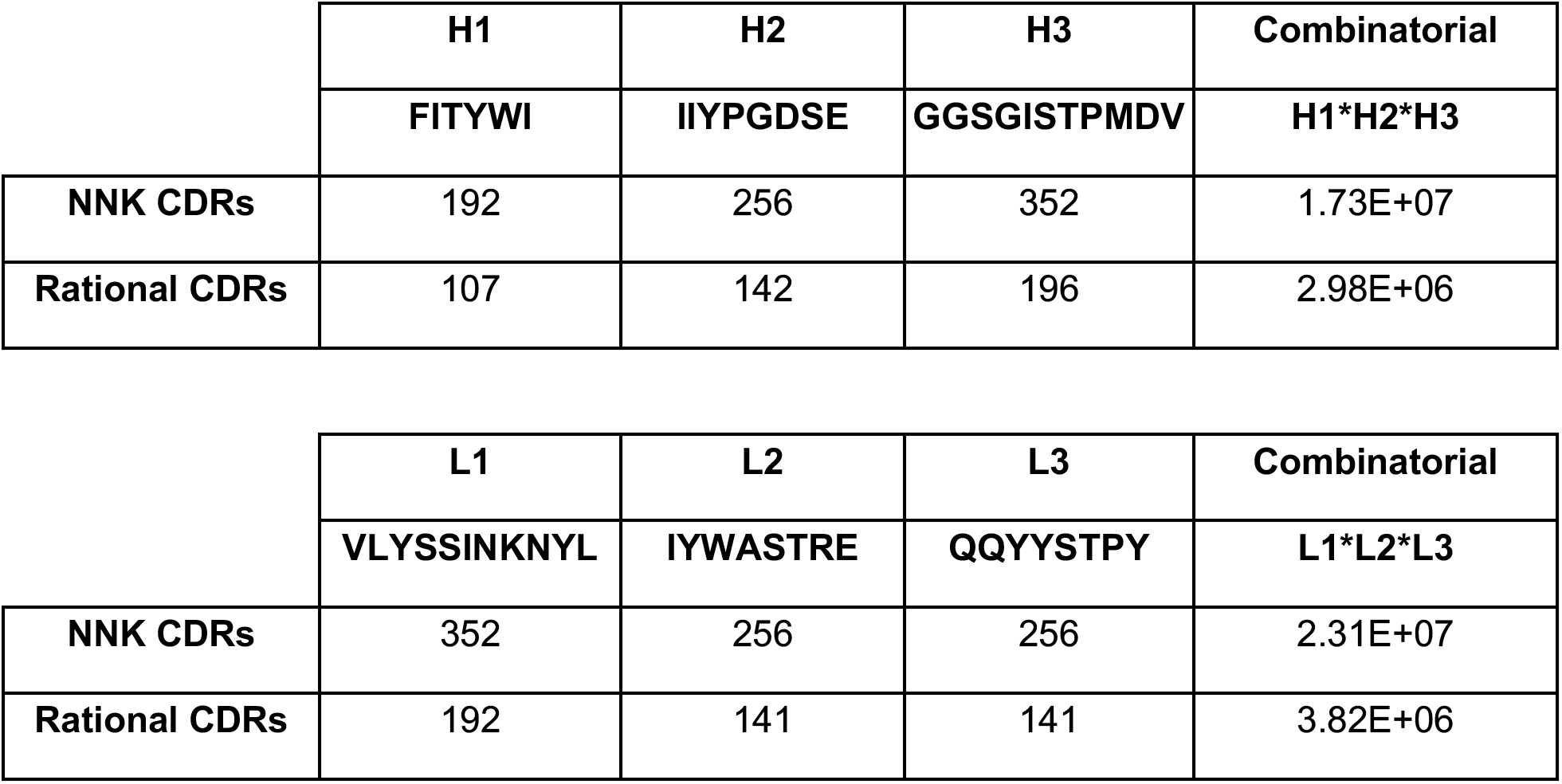
Theoretical library size. Related to Figure 1. Comparison of NNK generated libraries to rationally synthesized CDR libraries. The sequence of the starting CDR loop is given along with the number of variants in each loop that would result from NNK scanning or rational synthesis. The combinatorial size of the CDR1/2/3 library is given from the product of the 3 loops.

**Table S2.**
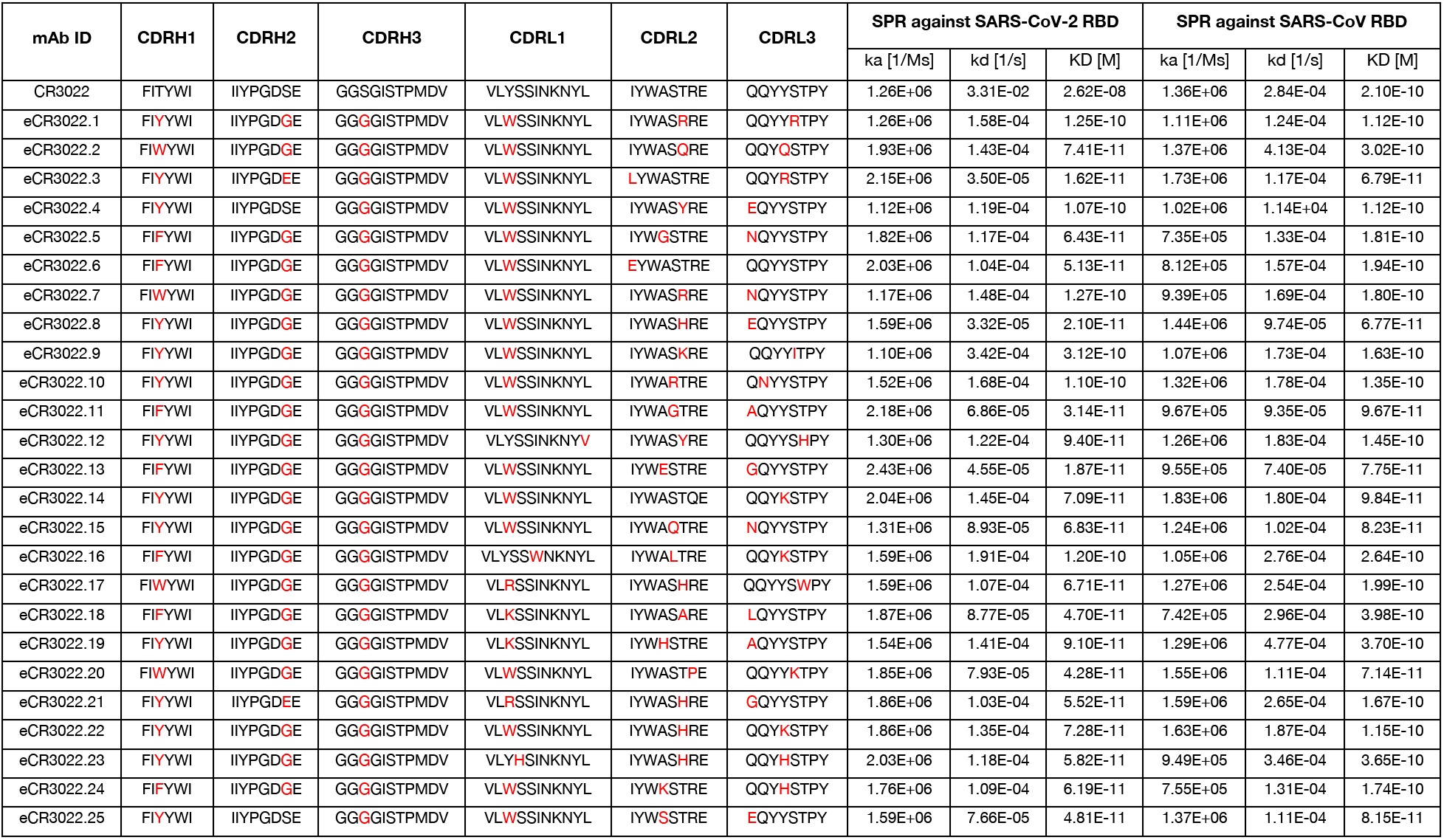
eCR3022 binding affinity and CDR loop sequences. Related to Figure 2. Summary table of parental CR3022 and 25 eCR3022 antibodies with sequences of 6 CDR loops and binding affinity against SARS-CoV-2 RBD and SARS-CoV-1 RBD. Mutations of eCR3022 at CDR loops were highlighted in red. Antibodies were captured via Fc-capture to an anti-human IgG Fc antibody and varying concentrations of SARS-CoV-2 or SARS-CoV-1 RBD were injected using a multi-cycle method. Association and dissociation rate constants calculated through a 1:1 Langmuir binding model using the BIAevaluation software.

**Table S3.**
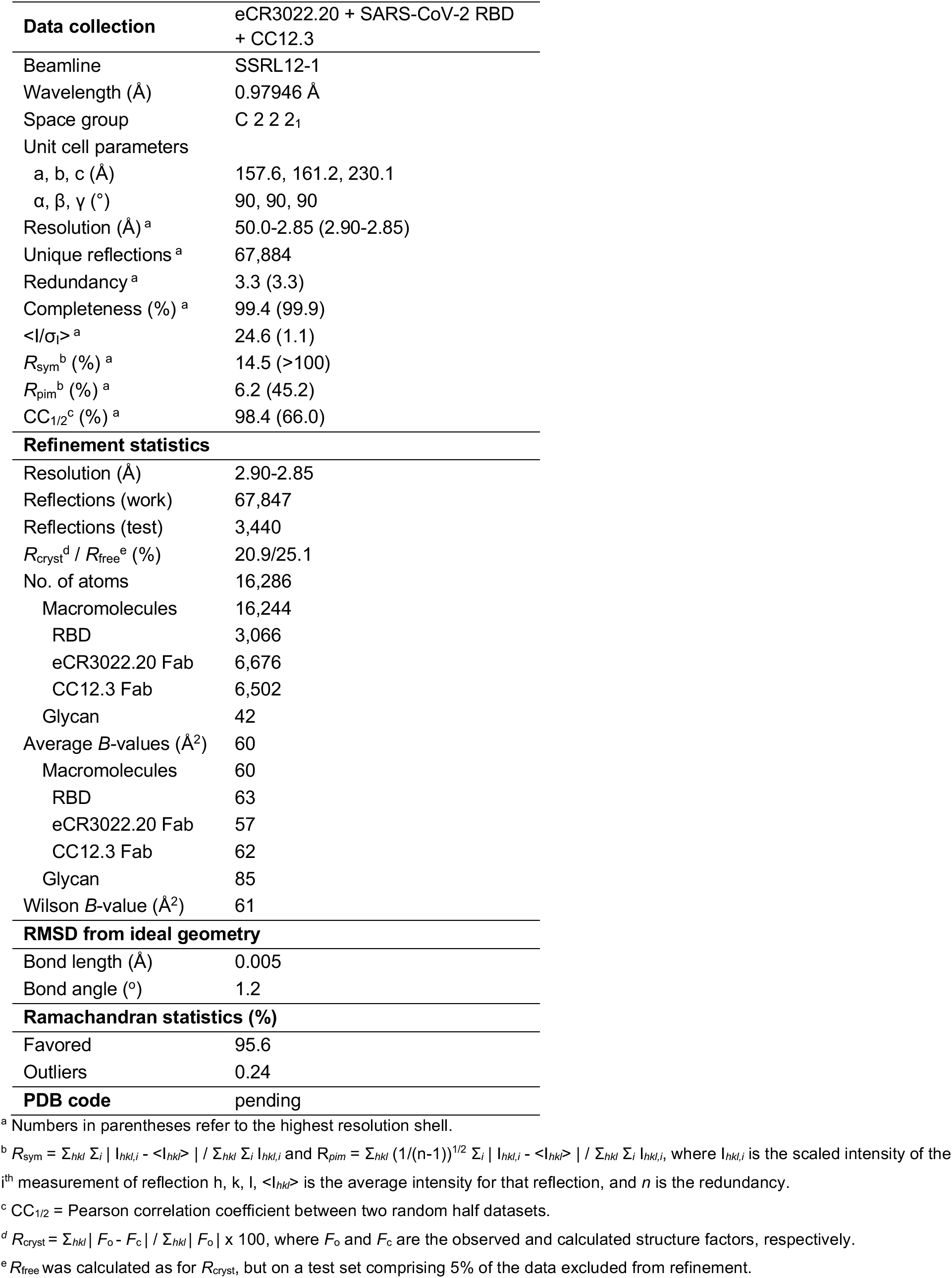
X-ray data collection and refinement statistics. Related to Figure 3.

**Table S4.**
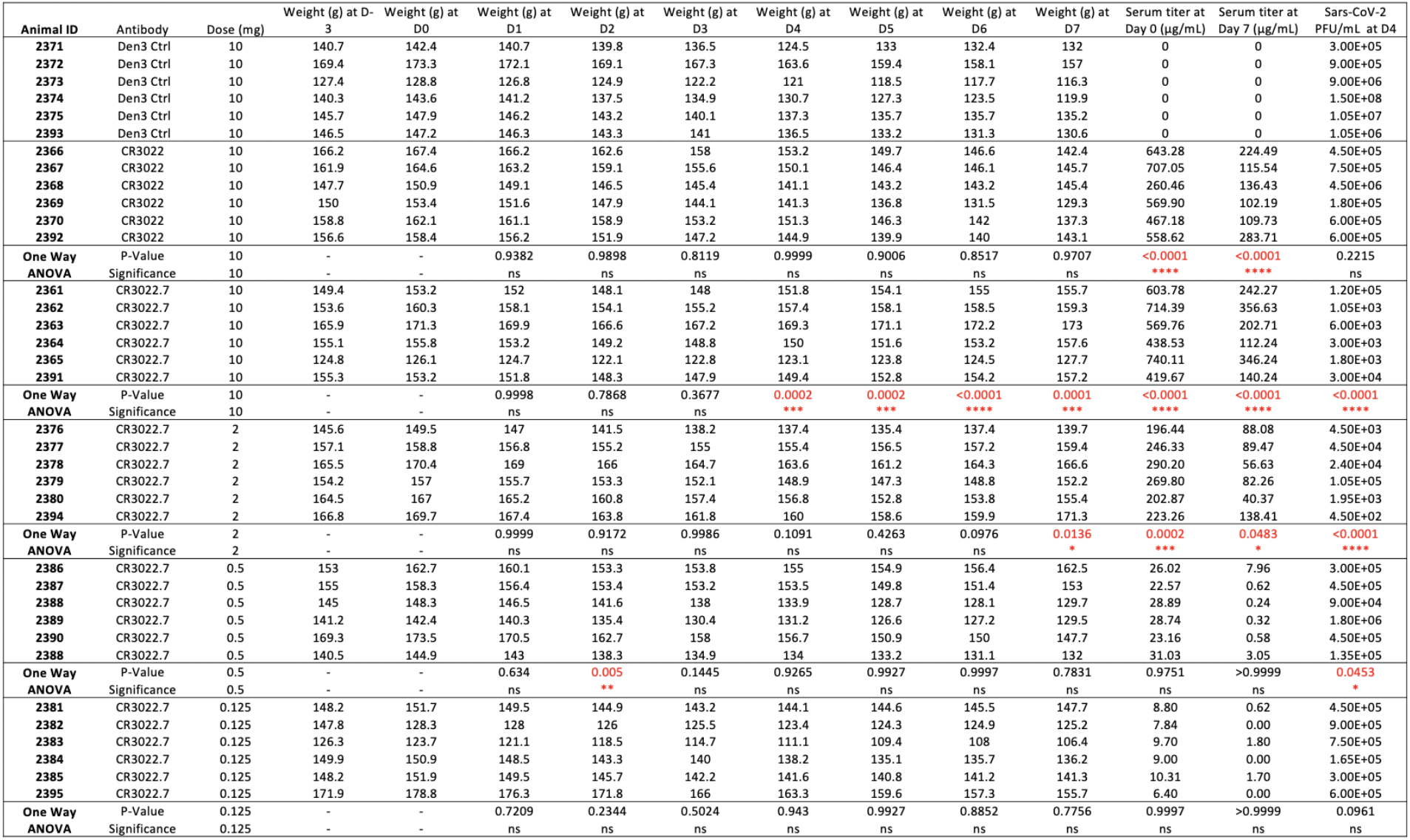
Hamster Protection Study Summary and Statistics. Related to Figure 4. Each group was compared to the Den3 IgG control group.

## SOM Figures

**Figure S1.**
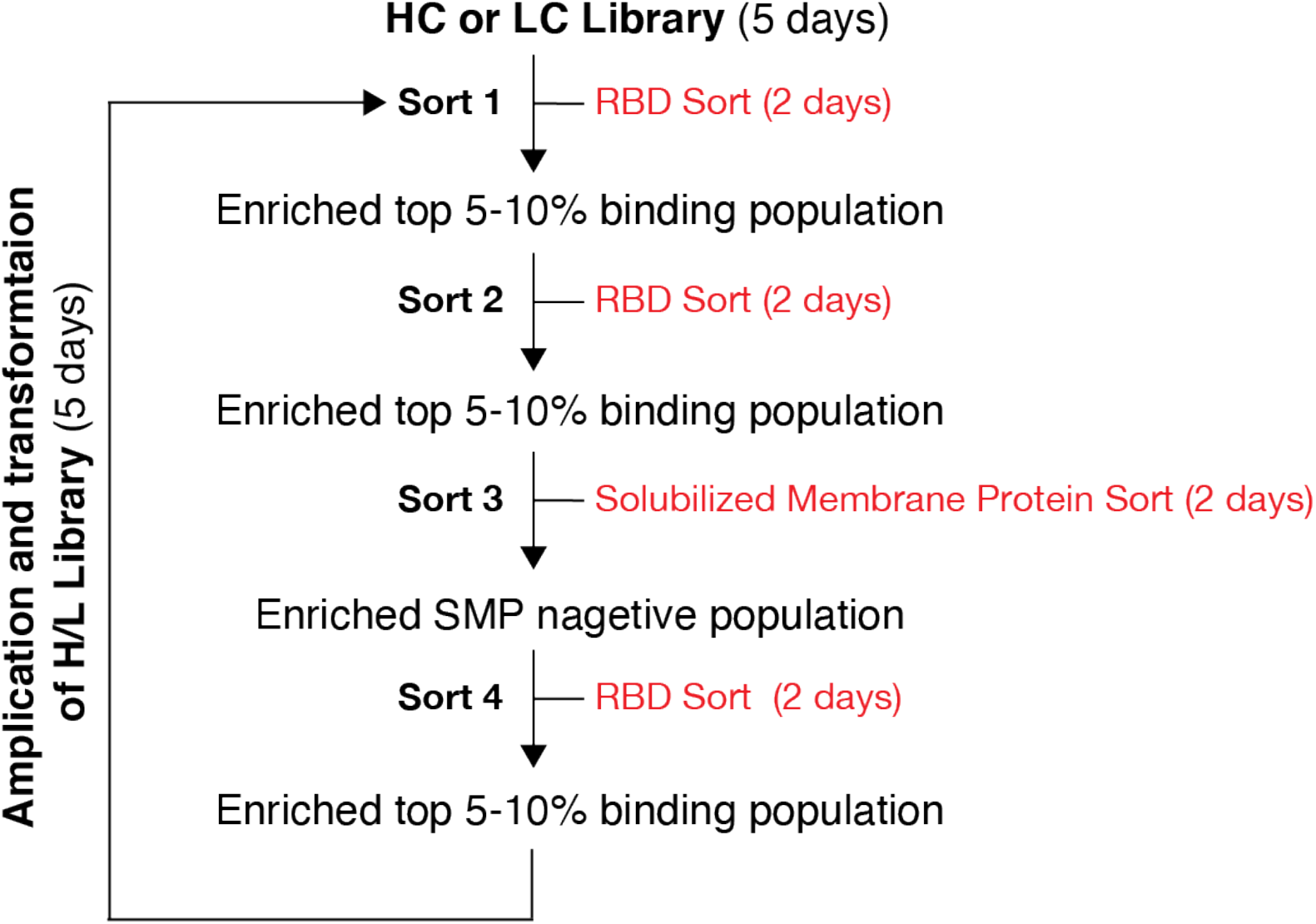
Overview of the library sorting process. Related to Figure 1. Schematic illustration of yeast population enrichment from HC, LC and combinatorial H/L libraries following four rounds of FACS selection.

**Figure S2.**
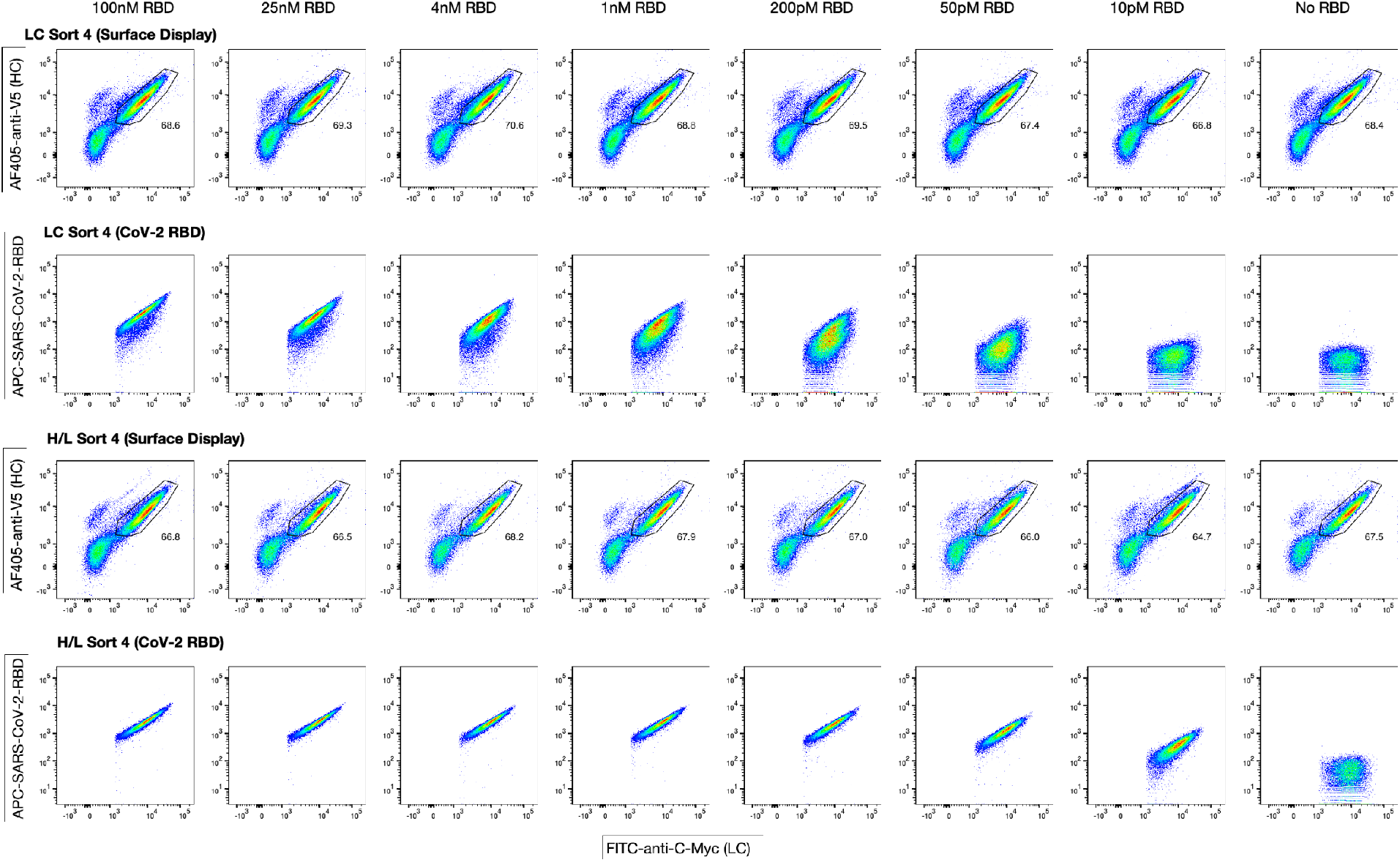
Representative FACS plots of CR3022 LC and H/L libraries in sort 4. Related to Figure 1. Yeast cells were induced and grown overnight at 30°C. Surface antibody display frequency was determined by staining with AF405-anti-V5 antibody (HC) and FITC-anti-c-Myc (LC). Cells were also labeled with different unsaturated concentrations of biotinylated SARS-CoV-2 RBD: 100 nM, 25 nM, 4 nM, 1 nM, 200 pM, 50 pM, 10 pM respectively. Labeled cells were further stained with APC conjugated streptavidin. FACS analysis was performed by BD FACSlyrics.

**Figure S3.**
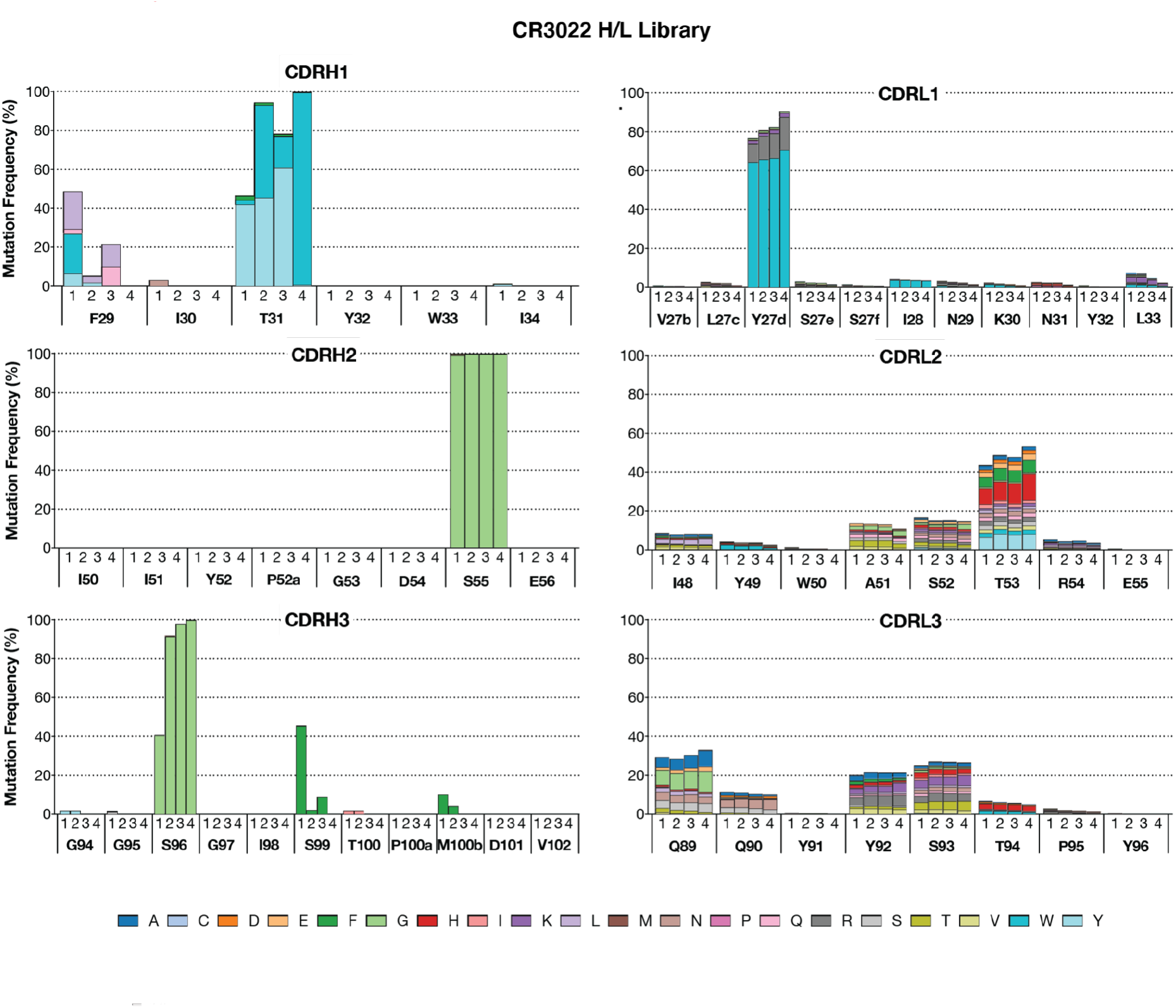
Deep sequencing analysis of mutations of CDR loops from the CR3022 combinatorial H/L library. Related to Figure 1. HC and LC from the combinatorial H/L library were amplified separately and then loaded onto Illumina Miseq sequencer using a Miseq Reagent V3 kit (600 cycles). Mutations in each CDR loop of HC (left) and LC (right) after each round of FACS sort were highlighted in colors corresponding to the key.

**Figure S4.**
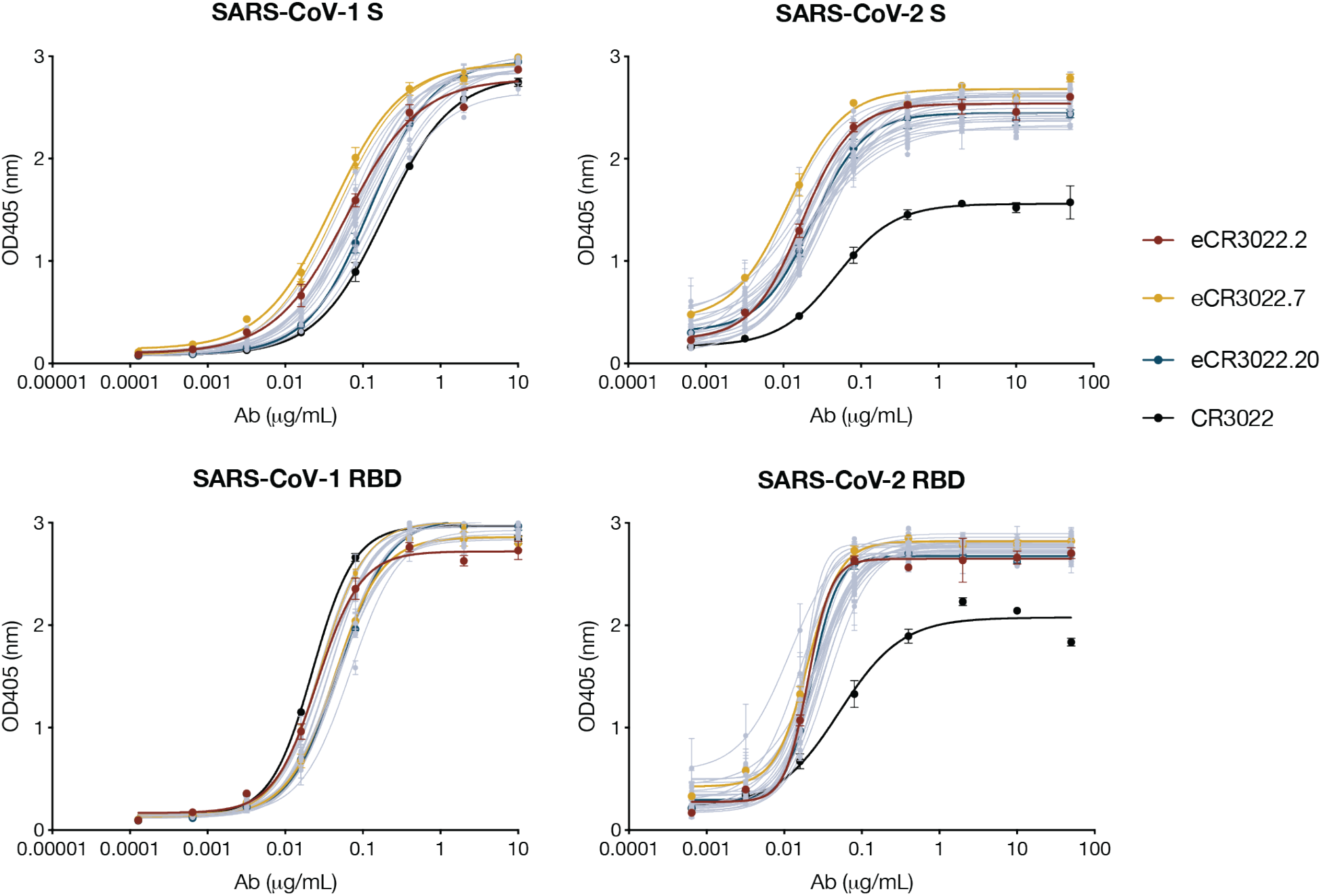
eCR3022 variants ELISA binding to SARS-CoV-1 and SARS-CoV-2 RBD and S proteins. Related to Figure 2. eCR3022 and parental CR3022 antibodies were evaluated binding against his-tagged SARS-CoV-1, SARS-CoV-2 RBD and S proteins. Each sample was tested in duplicates. Error bars represent standard deviations.

**Figure S5.**
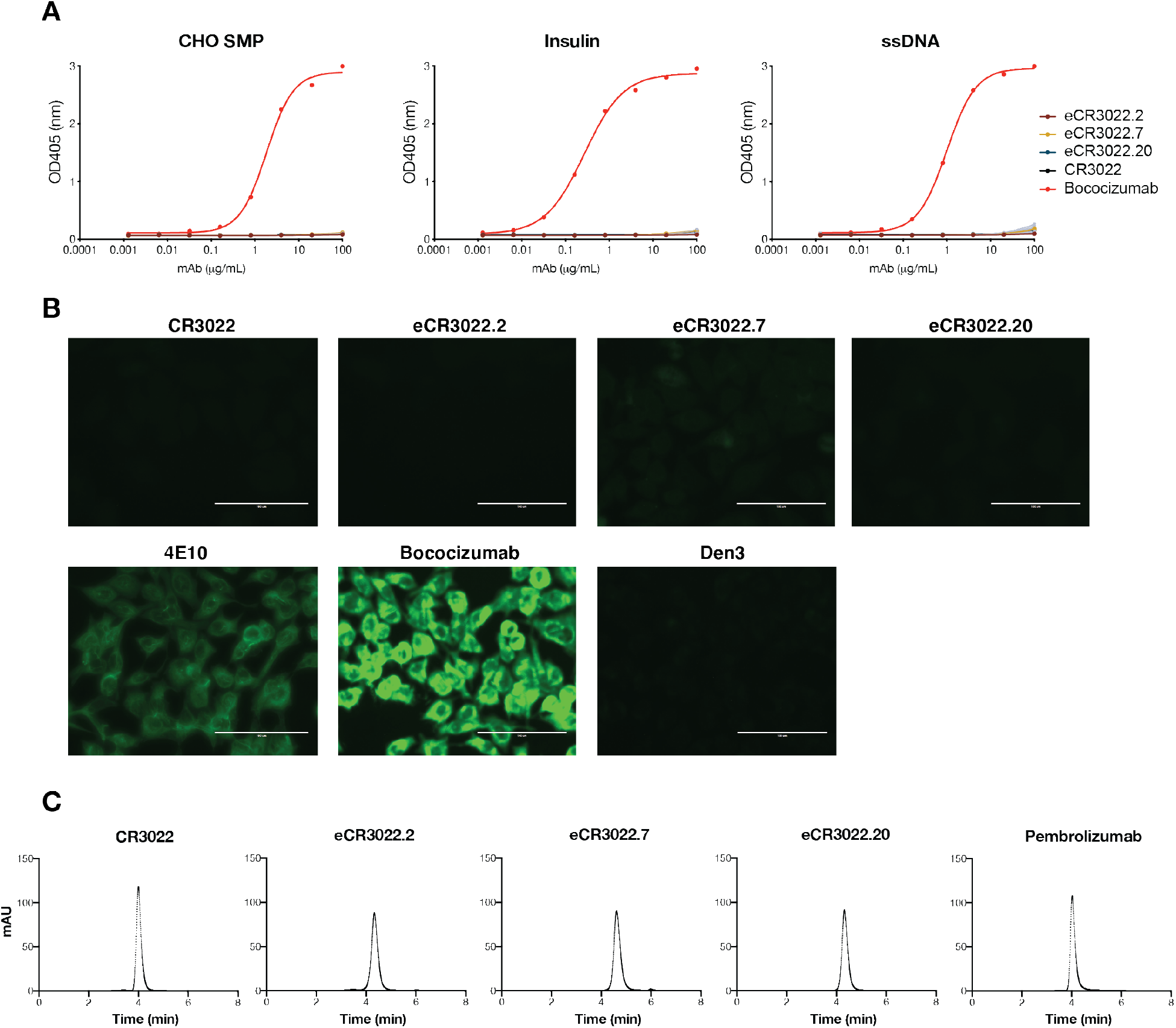
Evaluation of eCR3022 variants for polyreactivity and biopharmaceutical analysis. Related to Figure 2. Antibodies were tested by ELISA for binding against polyspecific reagents (PSR) against Chinese hamster ovary cells (CHO) solubilized membrane protein (SMP), insulin, single-strand DNA (ssDNA) (**A**) and by binding to immobilized HEp2 epithelial cells (**B**). Antibodies were further analyzed by Agilent size-exclusion chromatography (SEC) column (**C**) with an FDA-approved antibody Pembrolizumab as positive control.

**Figure S6.**
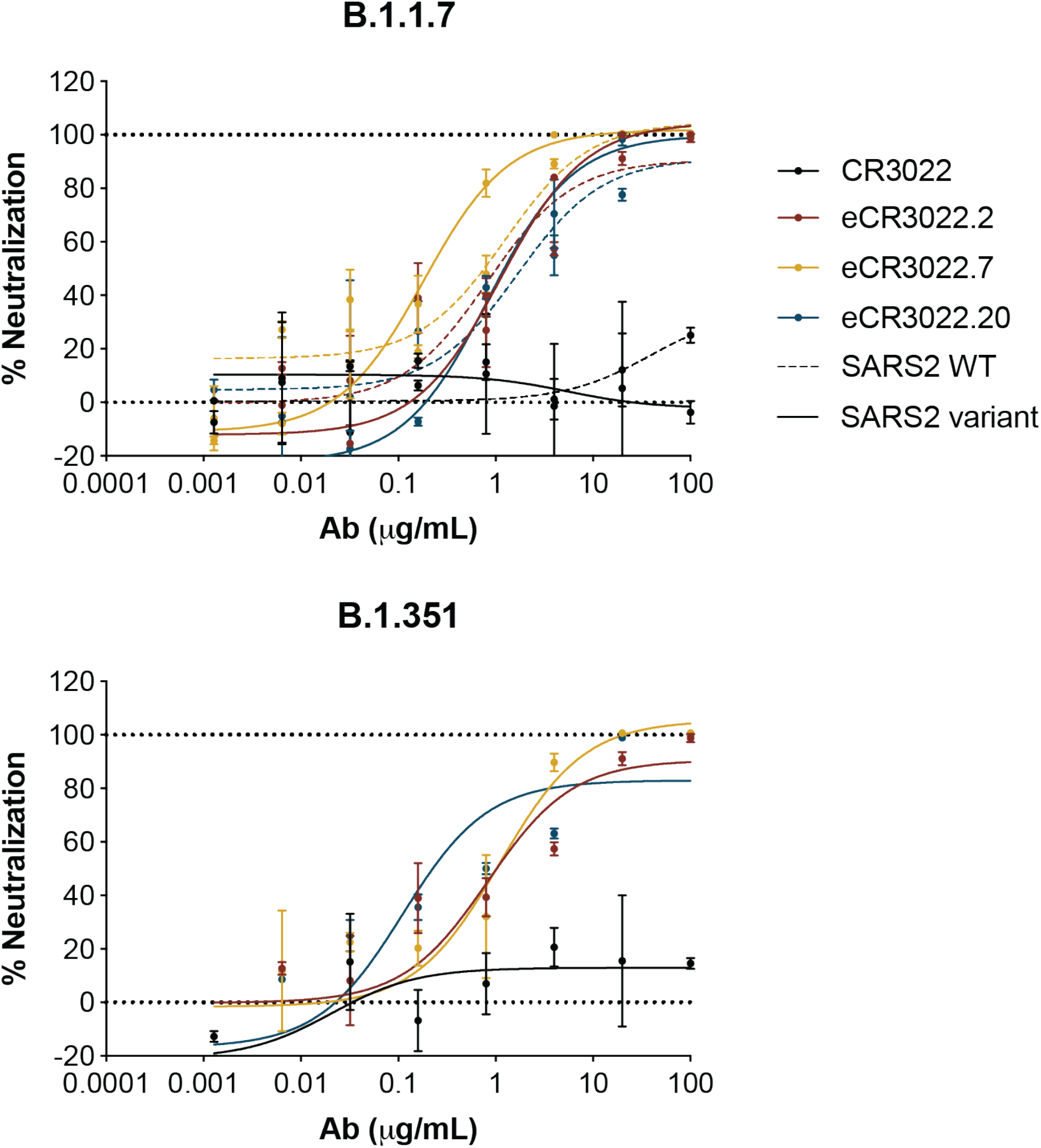
Representative neutralization of pseudotyped B.1.1.7 and B.1.351 variants. Related to Figure 2. Neutralization curves of parental CR3022 and eCR3022 antibodies against pseudotyped B.1.1.7 and B.1.351 lineages with full-spike mutations. Solid lines represent neutralization curves against SARS-CoV-2 variants while dashed lines represent curves against wildtype virus. Error bars represent standard deviations.

**Figure S7.**
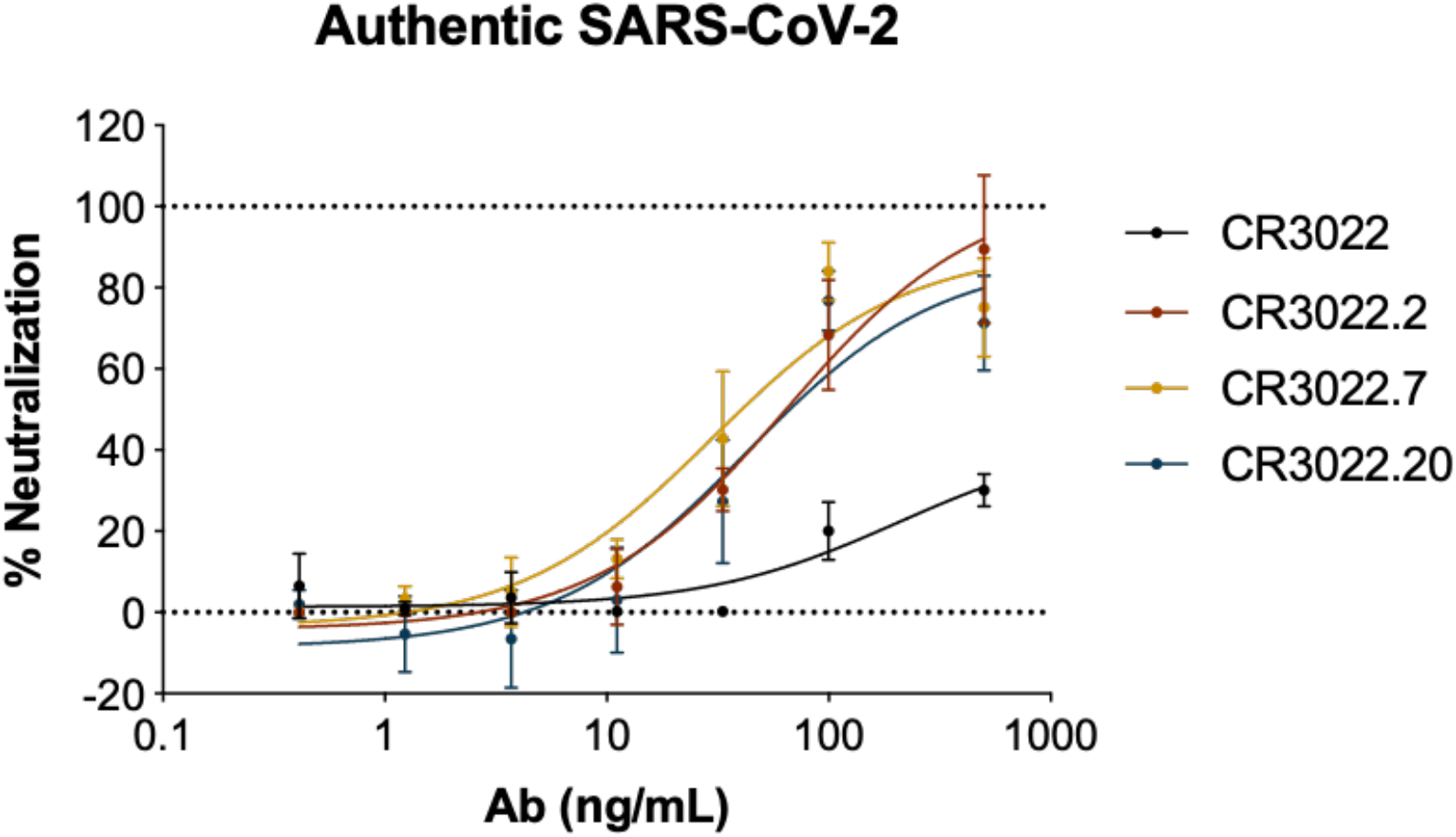
Representative neutralization of authentic SARS-CoV-2. Related to Figure 2. Neutralization curves of parental CR3022 and eCR3022 antibodies against authentic SARS-CoV-2 (USA-WA1/2020). Each sample was tested in duplicates. Error bars represent standard deviations.

**Figure S8.**
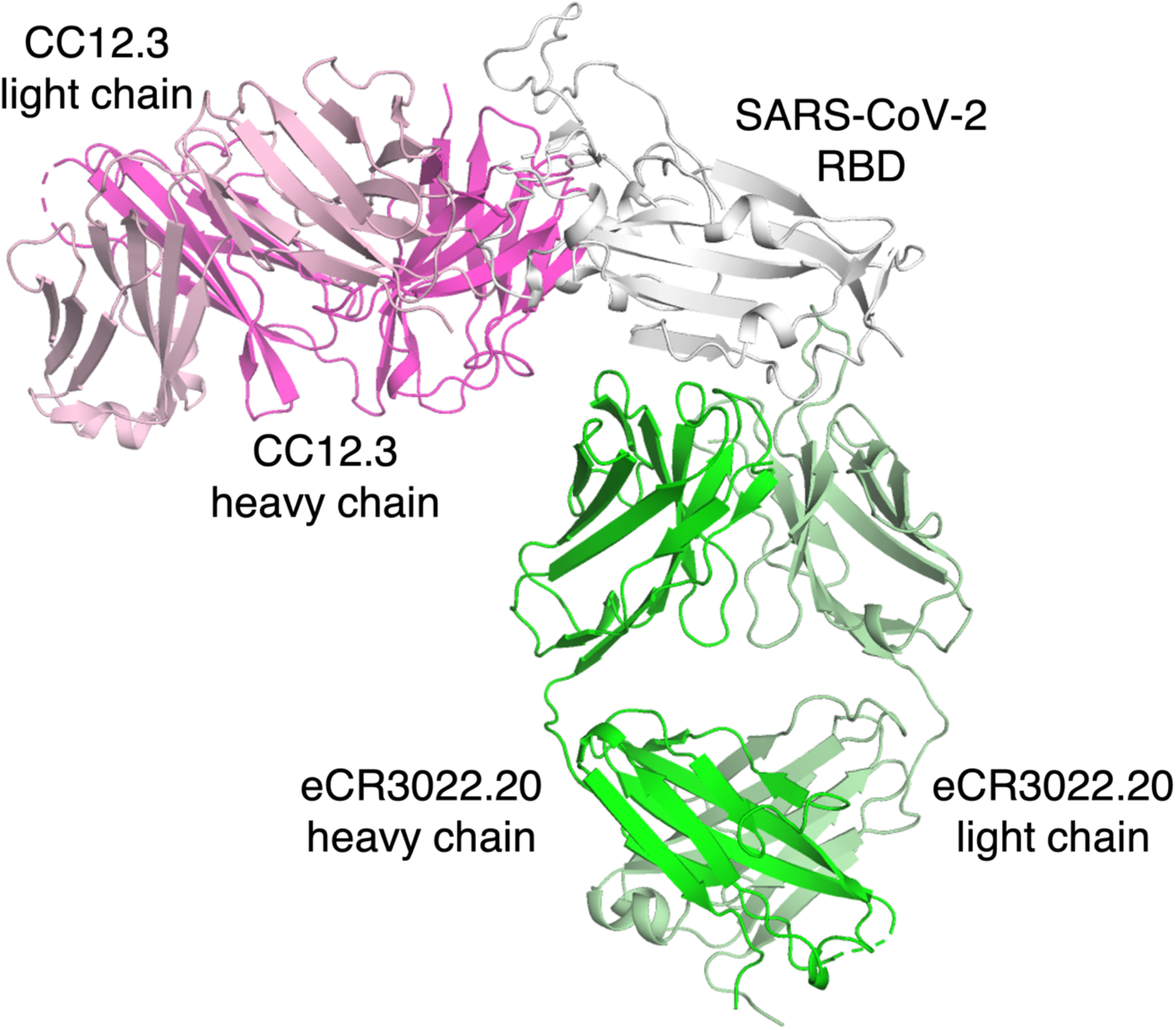
Crystal structure of SARS-CoV-2 RBD in complex with Fabs eCR3022.20 and CC12.3. Related to Figure 3. The binding site of eCR3022.20 (Fab heavy and light chains shown in green and light green, respectively) on the RBD (white) is distinct from that of CC12.3 (Fab heavy and light chains shown in magenta and light pink, respectively), which binds to the receptor binding site.

**Figure S9.**
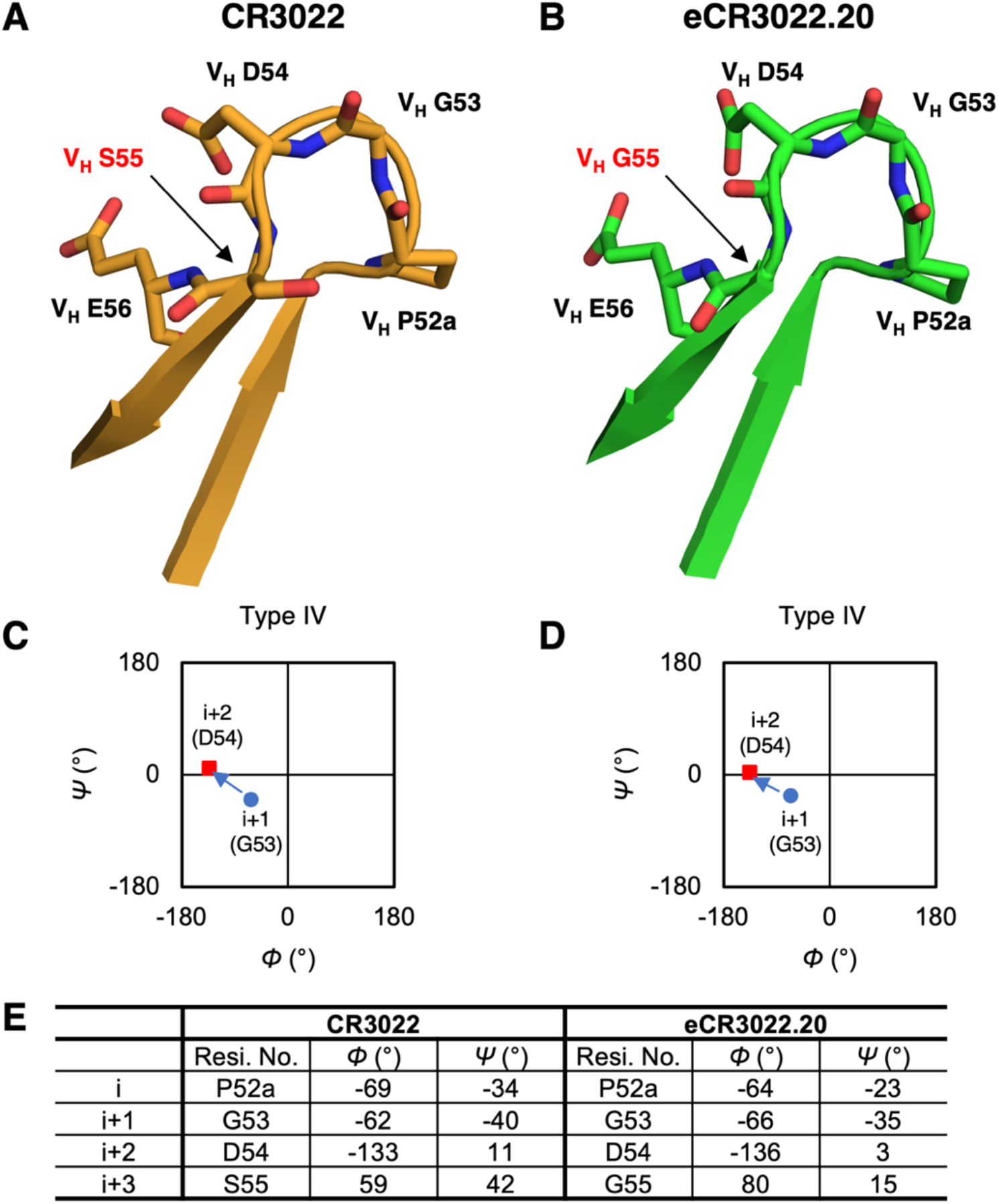
CDR H2 of CR3022 and eCR3022.20 form a type IV β turn. Related to Figure 3. (**A-B**) Comparison between the H2 CDRs of (**A**) CR3022 and (**B**) eCR3022.20. Residues that differ are highlighted in red letters. Our previous structure of CR3022 (PDB ID: 6W41) and the structure of eCR3022.20 are used for the comparison. (**C-D**) Ramachandran plots of the H2 CDRs of (**C**) CR3022 and (**D**) eCR3022.20 indicate type-IV β turns (V_H_ ^52a^PGDS^55^) for both H2 CDR loops are they deviate slightly from a type 1 β turn. Phi and psi angles of the residues i+1 (G53) and i+2 (D54) are shown as blue circles and red squares, respectively.

**Figure S10.**
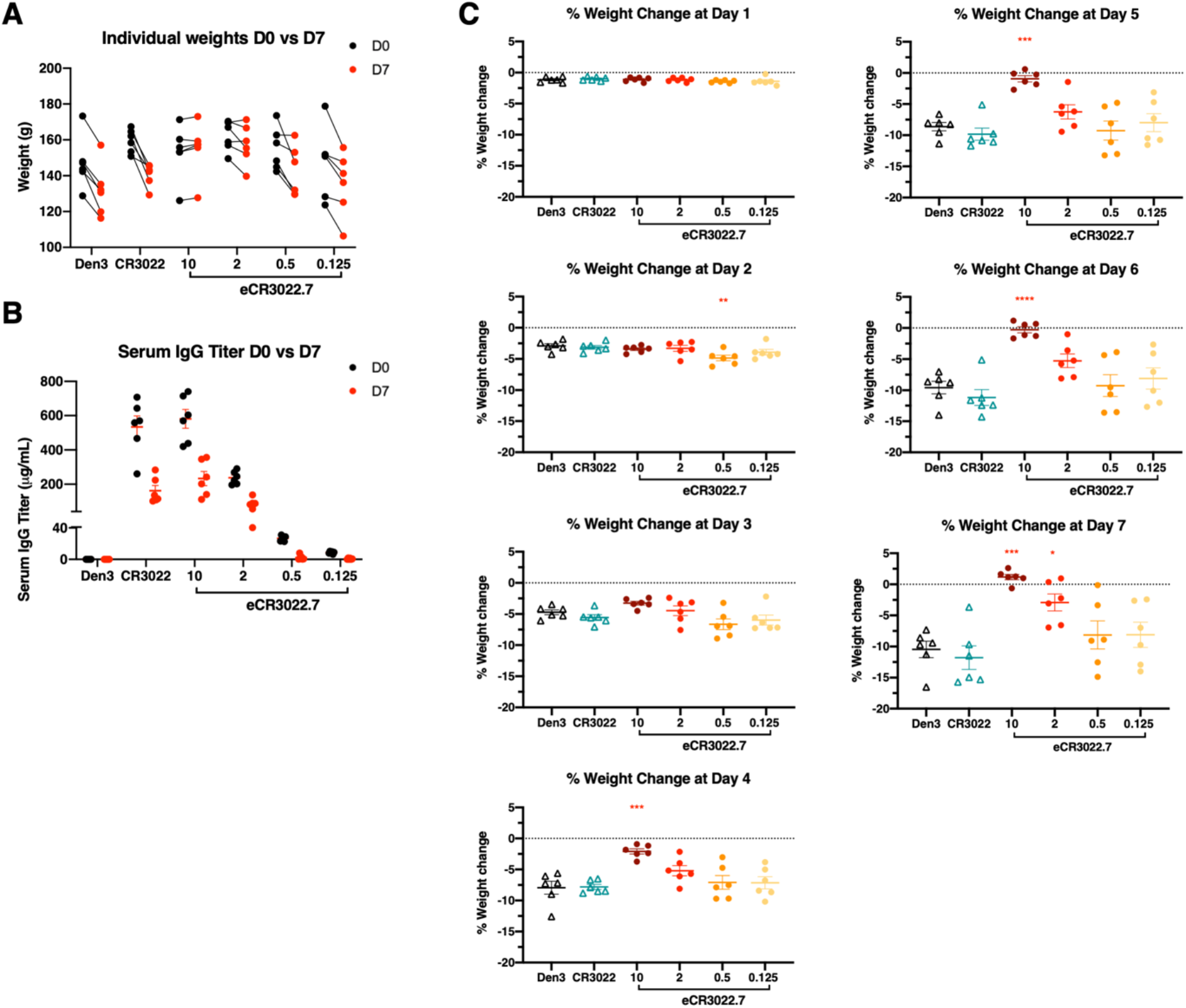
Animal protection studies. Related to Figure 4. (**A**) Weights of animals at time of challenge (Day 0) compared to weights at time of sacrifice (Day 7). (B) Serum human IgG concentration at time of infection (Day 0) compared to sacrifice (Day 7). (C) Percent weight loss by day compared to weights recorded at time of infection at day 0.

**Figure S11.**
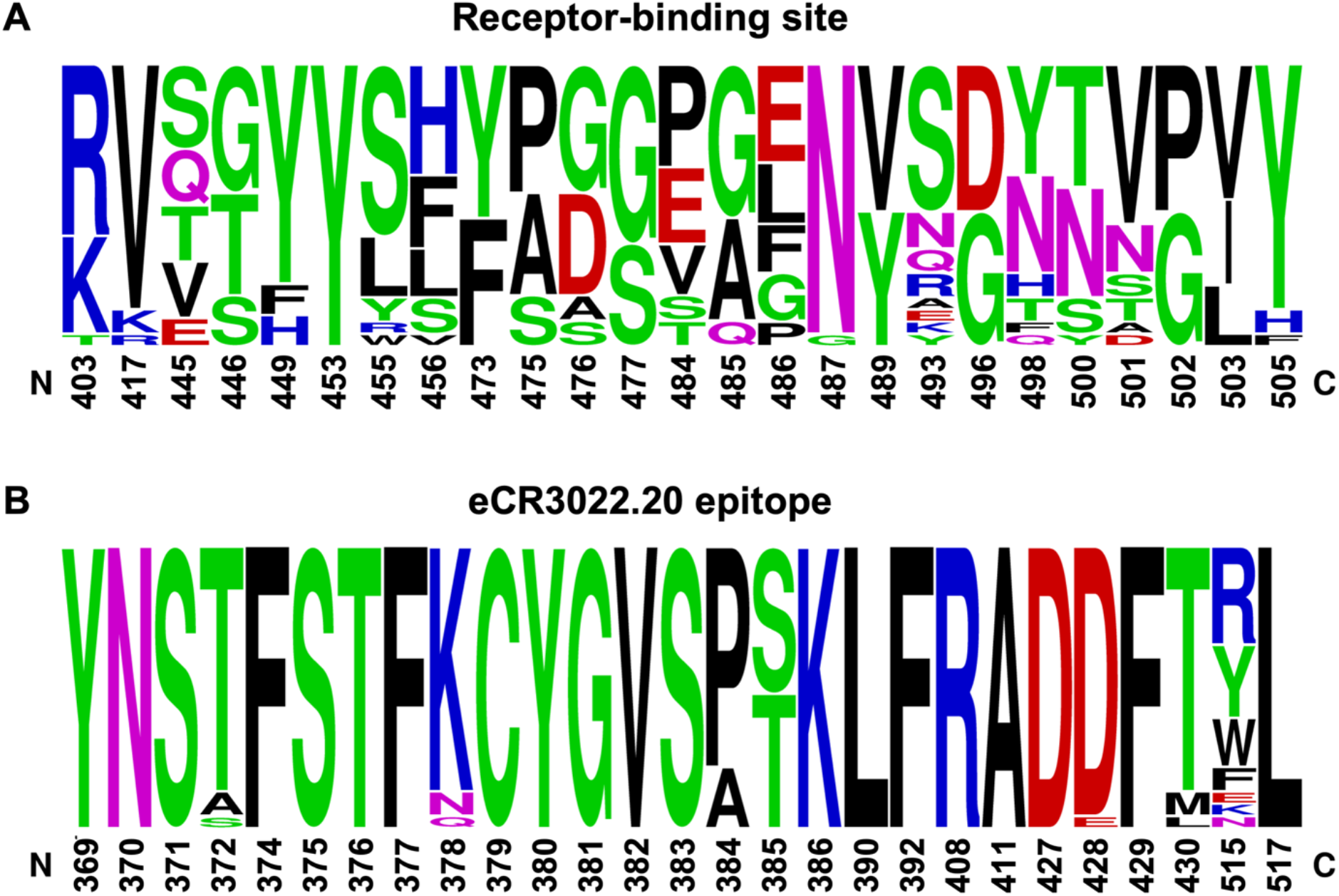
Sequence conservation of the eCR3022.20 epitope versus the SARS-CoV RBS. Residues that interact with **(A)** ACE2 and **(B)** eCR3022.20 (defined by BSA > 0 Å^2^) are shown as a sequence logo (Crooks et al., 2004). BSA values are calculated with PISA (Krissinel and Henrick, 2007) using the SARS-CoV-2 RBD/ACE2 structure (PDB 6M0J) (Lan et al., 2020) and the CR3022.20/RBD structure determined here. Sequences of 22 Sarbecoviruses including SARS-CoV-2, SARS-CoV and SARS-related coronaviruses (SARSr-CoVs) were used for this analysis: NCBI Reference Sequence YP_009724390.1 (SARS-CoV-2), GenBank ABF65836.1 (SARS-CoV), GenBank ALK02457.1 (Bat SARSr-CoV WIV16), GenBank AGZ48828.1 (Bat SARSr-CoV WIV1), GenBank ACU31032.1 (Bat SARSr-CoV Rs672), GenBank AIA62320.1 (Bat SARSr-CoV GX2013), GenBank AAZ67052.1 (Bat SARSr-CoV Rp3), GenBank AIA62300.1 (Bat SARSr-CoV SX2013), GenBank ABD75323.1 (Bat SARSr-CoV Rf1), GenBank AIA62310.1 (Bat SARSr-CoV HuB2013), GenBank AAY88866.1 (Bat SARSr-CoV HKU3-1), GenBank AID16716.1 (Bat SARSr-CoV Longquan-140), GenBank AVP78031.1 (Bat SARSr-CoV ZC45), GenBank AVP78042.1 (Bat SARSr-CoV ZXC21), GenBank QHR63300.2 (Bat CoV RaTG13), NCBI Reference Sequence YP_003858584.1 (Bat SARSr-CoV BM48-31), GISAID EPI_ISL_410721 (Pangolin-CoV Guandong2019), GenBank QIA48632.1 (Pangolin-CoV Guangxi), GenBank AGZ48806.1 (Bat SARSr-CoV RsSHC0144), GenBank ATO98120.1 (Bat SARSr-CoV Rs4081), GenBank AGC74176.1 (Bat SARSr-CoV Yun11), GenBank APO40579.1 (Bat SARSr-CoV BtKY72).

## MATERIALS AND METHODS

### Antibody library generation

CR3022 heavy chain and light chain affinity maturation libraries were synthesized by Twist Bioscience (San Francisco). Mutations were included in the CDR loops, based on the following definitions: CDRH1 = GYGFITYWI, CDRH2 = IIYPGDSET, CDRH3 = GGSGISTPMDV, CDRL1 = VLYSSINKNYL, CDRL2 = IYWASTRE, CDRL3 = QQYYSTPY. At each position in the CDR loop, mini libraries were synthesized that encoded all possible single mutations from the starting sequence, excluding variants where the substitution was to a cysteine or methionine and variants that created an N-linked glycosylation motif. The CDR1/2/3 mini-libraries were assembled into combinatorial heavy chain and light chain libraries.

The libraries were displayed on the surface of yeast as molecular Fab using the pYDSI vector, a yeast display vector containing the bidirectional Gal1-10 promoter that was based on the design of a previously described vector (Wang et al., 2018), omitting the leucine-zipper dimerization domains. The heavy chain contains a C-terminal V5 epitope tag and the light chain contains a C-terminal C-myc epitope tag to assess the amount of Fab displayed on the surface of the yeast. The HC library was generated by cloning the HC CDR1/2/3library into a vector already containing the invariant CR3022 light chain by homologous recombination, and the LC library was generated by doing the inverse. The HC/LC library was generated by amplifying the HC and LC sequences with primers overlapping in the Gal1-10 promoter. The recovered Gal-HC and Gal-LC fragments were ligated via Gibson assembly and amplified. The resulting LC-Gal1-10-HC product was cloned into empty pYDSI by homologous recombination.

### Yeast transformation

Yeast transformation was performed as described previously. In brief, the colony of *Saccharomyces cerevisiae* YVH10 cells (ATCC, MYA4940) was inoculated in 2mL YPD medium (Dissolve 20 g dextrose, 20 g peptone and 10 g yeast extract in deionized H_2_O to a volume of 1 liter and sterilize by filtration) and shaken overnight at 30°C. The overnight culture was expanded in 50 mL YPD medium and shaken at 30°C until the absorbance was around 1.5 at 600 nm. Yeast cells were spun down and resuspended with 25 mL of 100 mM lithium acetate. 250 μL of 1 M DTT was added to the cells and mixed rapidly. After shaking at 30°C for 10 min, cells were spun down and washed with 25 mL of pre-chilled deionized H_2_O. After centrifuge, the cell pellet was suspended with pre-chilled deionized H_2_O to a final volume of 500 μL. After this step, yeast cells were electrocompetent and ready for transformation.

1 μg of linearized vector DNA was mixed with 5 ug of insert library DNA in an Eppendorf tube on ice. 250 μL of electrocompetent cells were transferred to the tube and incubated for 10 min on ice. Then the cells and DNA mixture were transferred to cuvette and inserted into Gene Pulser Xcell Electroporation System (Biorad) using following settings:

Square wave
Voltage = 500 V
Pulse length = 15.0 ms
\# pulses = 1
Pulse interval = 0
Cuvette = 2 mm

After electrophoresis, cells were shocked by immediately adding 1 mL of pre-warmed YPD medium. Cells were then transferred to a 50 mL tube for outgrowth. After shaking 200 rpm at 30°C for 1 h, cells were centrifuged and resuspended in synthetic drop-out medium without tryptophan (with 1 % Penicillin/Streptomycin) and shaken overnight to grow.

### Yeast library labeling and sorting

After yeast transformation, yeast cells were expanded and split once for better display efficiency. Before staining, cells were induced overnight at 30°C by SGCAA induction medium (dissolve 20 g galactose, 1 g glucose, 6.7 g yeast nitrogen base without amino acid, 5 g bacto casamino acids, 5.4 g Na_2_HPO_4_, 8.56 g NaH_2_PO_4_·H_2_O, 8.56 mg uracil to 1 L deionized water, pH 6.5, and sterilize by filtration). For each library, in the first round of selection, 5 x 10^7^ of yeast cells were stained per sample. In the second to final round of selection, 1 x 10^7^ cells were stained. Yeast cells were firstly spun down and washed with PBS/1% BSA, then incubated with biotinylated SARS-CoV-2 RBD or S or HEK cell membrane protein at several non-depleting concentrations respectively for at least 30 min at 4°C. After washing, yeast cells were stained with FITC-conjugated chicken anti-C-Myc antibody (Immunology Consultants Laboratory, CMYC-45F), AF405-conjugated anti-V5 antibody (made in house), and streptavidin-APC (Invitrogen, SA1005) in 1:100 dilution for 20 min at 4 °C. After washing, yeast cells were resuspended in 1 mL of PBS/1% BSA and loaded on BD FACSMelody cell sorter. Top 5-10% of cells with high binding activity to a certain SARS-CoV-2 RBD labeling concentration were sorted and spun down. Sorted cells were expanded in 2 mL of synthetic drop-out medium without tryptophan (Sigma-Aldrich, Y1876-20G) supplemented with 1% Penicillin/Streptomycin (Corning, 30-002-C) at 30°C overnight.

### Deep sequencing and analysis

After each sort, a fragment of cell population was expanded in 2 mL of synthetic drop-out medium without tryptophan supplemented with 1% Penicillin/Streptomycin overnight at 30°C. Yeast cells were then spun down, cell pellet was resuspended with 250 μL of buffer P1 (with RNAse added) (Qiagen, 27104) by pipetting up and down. 5 μL of Zymolyase (Zymo Research, E1005) was added to digest yeast cell walls and incubated at 37°C for 1 h. Cells were then lysed, neutralized, and DNA was purified according to manufacturer’s instructions (Qiagen, 27104). After that, CR3022 HC and LC fragments from post-sorted plasmid DNA were amplified by following CR3022-HC and CR3022-LC primer mixture respectively.

CR3022.HC-NGSFa:GTCTCGTGGGCTCGGAGATGTGTATAAGAGACAGGAGTCTCTGAAGATCTCCTGT AAGGG;

CR3022.HC-NGSFb:GTCTCGTGGGCTCGGAGATGTGTATAAGAGACAGHHGAGTCTCTGAAGATCTCCT GTAAGGG;

CR3022.HC-NGSFc:GTCTCGTGGGCTCGGAGATGTGTATAAGAGACAGHHHHGAGTCTCTGAAGATCTC CTGTAAGGG;

CR3022.HC-NGSRa:TCGTCGGCAGCGTCAGATGTGTATAAGAGACAGGAGACGGTGACCGTGGTTC;

CR3022.HC-NGSRb:TCGTCGGCAGCGTCAGATGTGTATAAGAGACAGHHGAGACGGTGACCGTGGTTC;

CR3022.HC-NGSRc:TCGTCGGCAGCGTCAGATGTGTATAAGAGACAGHHHHGAGACGGTGACCGTGGT TC;

CR3022.LC-NGSFa:GTCTCGTGGGCTCGGAGATGTGTATAAGAGACAGGGAGAAAGAGCCACCATCAACTG;

CR3022.LC-NGSFb:GTCTCGTGGGCTCGGAGATGTGTATAAGAGACAGHHGGAGAAAGAGCCACCATCA ACTG;

CR3022.LC-NGSFc:GTCTCGTGGGCTCGGAGATGTGTATAAGAGACAGHHHHGGAGAAAGAGCCACCAT CAACTG;

CR3022.LC-NGSRa:TCGTCGGCAGCGTCAGATGTGTATAAGAGACAGGATCTCCACCTTGGTCCCTTG;

CR3022.LC-NGSRb:TCGTCGGCAGCGTCAGATGTGTATAAGAGACAGHHGATCTCCACCTTGGTCCCTT G;

CR3022.LC-NGSRc:TCGTCGGCAGCGTCAGATGTGTATAAGAGACAGHHHHGATCTCCACCTTGGTCCC TTG.

1μL of primer mixture (10 μM) were used to amplify the HC and LC DNA respectively after every round of FACS selection with 2 μL of post-sorted plasmid DNA, 10 μL of 5X Phusion HF buffer, 35.5 μL of H_2_O, and 0.5 μL of Phusion enzyme (ThermoFisher, F530L) using the following PCR program: 1min at 98°C; 32 cycles of 10 s at 98°C, 15 s at 65°C, 30s at 72°C; followed by 5 min at 72°C. After PCR clean up, second round of PCR was performed by adding 2 μL of first round PCR product, 5 μL of 4 μM barcode nextera adapter primer mixture, 0.4 uL of 10mM dNTP, 4 uL of 5X Phusion HF buffer (ThermoFisher, F518L), 8.4 μL of H_2_O, and 0.2 μL of Phusion enzyme and using the following PCR program: 1 min at 98°C; 7 cycles of 15 s at 98°C, 15 s at 68°C, 30 s at 72°C; followed by 5 min at 72°C. After PCR clean up, all the PCR products were combined and diluted to 15 pM in the final volume of 20 μL. To denature the DNA, 5 μL of the diluted library and 5 μL of freshly-prepared 0.2N NaOH were mixed and incubated at room temperature for 5 min. Then 990 μL of pre-chilled HyB buffer was added and mixed well. 570 μL of denatured library DNA and 30 μL of denatured PhiX control library (Illumina, FC-110-3001) were mixed, added into Miseq Reagent V3 kit (Illumina, MS-102-3003), and finally loaded onto Illumina Miseq Next Generation sequencer.

Paired FASTQs were checked for sequence quality using the FastQC package (FastQC v0.11.9). The forward and reverse reads were merged using BBMerge (version 38.87) from the BBTools suite using standard parameters (Bushnell et al., 2017). Merged reads with full sequence identity were clustered using VSEARCH (v2.15.1) (Rognes et al., 2016). Clustering was done using the “cluster_fast” method and fasta files were written including cluster abundance in the fasta header.

A custom python (Python 3.7) script was written to parse the vsearch output, translate the DNA sequences to amino acid sequences and count the CDR1, CDR2, and CDR3 positions. Using VSEARCH for clustering prior to translating and parsing the sequences improved performance substantially.

### Pacbio sequencing

Long Amp Taq Polymerase (New England Biolabs) was used to PCR amplify Plasmid DNA after sort 4 according to manufacturer’s protocol with the following primers:

CR3022_PCR1_FWD: /5AmMC6/GCAGTCGAACATGTAGCTGACTCAGGTCACCAAACAACAGAAGCAGTCCCA
CR3022_PCR1_REV: /5AmMC6/TGGATCACTTGTGCAAGCATCACATCGTAGGGAGTTCAGGTGCTGGTGAT.

First round PCR products were purified with SPRI beads (Beckman Coulter) and 10 uL of purified PCR product was used in a second round of index PCR with the following primers:

bc_1004_FWD_PacB_Univ.PCR: GGGTCACGCACACACGCGCGgcagtcgaacatgtagctgactcaggtcac
bc_1028_REV_PacB_Univ.PCR: CAGTGAGAGTCAGAGCAGAGtggatcacttgtgcaagcatcacatcgtag

DNA sample was again purified with SPRI beads, then submitted to GeneWiz, where a PacBio SMRTbell amplicon library was prepared per the manufacturer’s protocol and sequenced on the PacBio Sequel platform with v3.0 chemistry. The generated subreads were demultiplexed and circular consensus sequence (CCS) reads were obtained using the CCS algorithm within PacBio ccs v4.2.0. The algorithm was run using the default parameters. A custom python (Python 3.7) script was written to parse the CCS fastq output, translate the DNA sequences to amino acid sequences and count the CDR1, CDR2, and CDR3 positions.

Sequence data that support the findings in this study are available at the NCBI Sequencing Read Archive (www.ncbi.nlm.nih.gov/sra) under BioProject number PRJNAXXXXXX. Python code will be available on github.

### Recombinant S and RBD production

SARS-CoV-1 (Genbank AAP13567) or SARS-CoV-2 (Genbank MN908947) S and RBD proteins were transiently expressed in Freestyle 293F system (ThermoFisher). In brief, S or RBD expression plasmids were cotransfected with 40K PEI (1 mg/mL) at a ratio of 1:3. After incubation for 30 min at RT, transfection mixture was added to Freestyle 293F cells at a density of approximately 1 million cells/mL. After 5 days, supernatants were harvested and filtered with a 0.22 μm membrane.The His-tagged proteins were purified with the HisPur Ni-NTA Resin (Thermo Fisher, 88222). After three columns of washing with 25 mM Imidazole (pH 7.4), proteins were eluted with an elution buffer (250 mM Imidazole, pH 7.4) at slow gravity speed (~4 sec/drop). Eluted proteins were buffer exchanged and concentrated with PBS using Amicon tubes (Millipore). The proteins were further purified by size exclusion chromatography (SEC) using Superdex 200 (GE Healthcare). The selected fractions were pooled and concentrated again for further use.

### Antibody production and purification

Monoclonal antibody was transiently expressed in the Expi293 system (ThermoFisher, A14635). In brief, antibody HC and LC plasmids were co-transfected at a ratio of 1:2.5 with transfection reagent FectoPRO (Polyplus 116-010). After 24 h of transfection, 300 mM of sterile sodium valproic acid solution (Sigma-Aldrich, P4543) and 45% D-(+)- glucose solution (Sigma Aldrich, G8769-100ML) were added to feed cells. After 4-5 days of transfection, supernatants were collected, sterile-filtered (0.22 μm), and IgG was purified with Protein A sepharose beads (GE Healthcare 17-5280-04).

### Pseudovirus neutralization assay

Pseudovirus was generated as described previously^2^. In brief, 12.5 μg of MLV gag/pol backbone (Addgene, 14887), 10 μg of MLV-CMV-Luciferase plasmid, and 2.5 μg of SARS-CoV-2-d18 spike plasmid were incubated with transfection reagent Lipofectamine 2000 (Thermo Fisher, 11668027) following manufacturer’s instructions for 20 min at RT. Then the mixture was transferred onto HEK 293T cells (ATCC, CRL-3216) in a 10 cm^2^ culture dish (Corning, 430293). After 12-16 h of transfection, culture medium was gently removed, fresh DMEM medium was added onto the culture dish. Supernatants containing pseudovirus were harvested after 48 h post transfection and frozen at −80 °C for long term storage.

In the neutralization assay, antibody samples were serially diluted with complete DMEM medium (Corning, 15-013-CV) containing 10% FBS (Omega Scientific, FB-02), 2 mM L-Glutamine (Corning, 25-005-Cl), and 100 U/mL of Penicillin/Streptomycin (Corning, 30-002-C). 25 μL/well of diluted samples were then incubated with 25 μL/well of pseudotyped virus for 1 h at 37 °C in 96-well half-area plates (Corning, 3688). After that, 50 μL of Hela-hACE2 cells were added at 10,000 cells/well onto each well of the plates. After 48 h of incubation, cell culture medium was removed, luciferase lysis buffer (25 mM Gly-Gly pH 7.8, 15 mM MgSO4, 4 mM EGTA, 1% Triton X-100) was added onto cells. Luciferase activity was measured by BrightGlo substrate (Promega, PR-E2620) according to the manufacturer’s instructions. mAbs were tested in duplicate wells and independently repeated at least twice. Neutralization IC_50_ values were calculated using “One-Site Fit LogIC50” regression in GraphPad Prism 8.0.

### Authentic SARS-CoV-2 neutralization assay

Vero E6 cells were seeded in 96-well half-well plates at approximately 8000 cells/well in a total volume of 50 μL complete DMEM medium the day prior to the addition antibody and virus mixture. The virus (500 plaque forming units/well) and antibodies were mixed, incubated for 30 minutes and added to the cells. The transduced cells were incubated at 37°C for 24 hours. Each treatment was tested in duplicate. The medium was removed and disposed of appropriately. Cells were fixed by immersing the plate into 4% formaldehyde for 1 hour before washing 3 times with phosphate buffered saline (PBS). The plate was then either stored at 4°C or gently shaken for 30 minutes with 100 μL/well of permeabilization buffer (PBS with 1% Triton-X). All solutions were removed, then 100 μl of 3% bovine serum albumin (BSA) was added, followed by room temperature (RT) incubation at 2 hours.

Primary antibodies against the spike protein were generated from a high-throughput process that screened a convalescent, coronavirus disease 2019 cohort (CC)^2^. A mix of primary antibodies consisting of CC6.29, CC6.33, CC6.36, CC12.23, CC12.25, in a 1:1 ratio, were used next. The primary antibody mixture was diluted in PBS/1% BSA to a final concentration of 2 μg/ml. The blocking solution was removed and washed thoroughly with wash buffer (PBS with 0.1% Tween-20). The primary antibody mixture, 50 μl/well, was incubated with the cells for 2 hours at RT. The plates were washed 3 times with wash buffer.

Peroxidase AffiniPure Goat Anti-Human IgG (H+L) secondary antibody (Jackson ImmunoResearch, 109-035-088) diluted to 0.5 μg/mLl in PBS/1% BSA was added at 50 μL/well and incubated for 2 hours at RT. The plates were washed 6 times with wash buffer. HRP substrate (Roche, 11582950001) was freshly prepared as follows: Solution A was added to Solution B in a 100:1 ratio and stirred for 15 minutes at RT. The substrate was added at 50 μL/well and chemiluminescence was measured in a microplate luminescence reader (BioTek, Synergy 2).

The following method was used to calculate the percentage neutralization of SARS-CoV-2. First, we plotted a standard curve of serially diluted virus (3000, 1000, 333, 111, 37, 12, 4, 1 PFU) versus RLU using four-parameter logistic regression (GraphPad Prism 8.0) below:

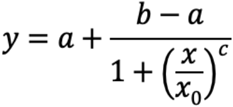

(y: RLU, x: PFU, a,b,c and x0 are parameters fitted by standard curve)

To convert sample RLU into PFU, use the equation below: (if y < a then x = 0)

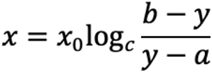

Percentage neutralization was calculated by the following equation:

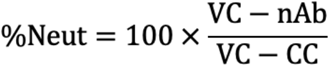

VC = Average of vehicle-treated control; CC = Average of cell only control, nAb, neutralizing antibody. PFU value was used for each variable indicated.

To compute neutralization IC_50_, logistic regression (sigmoidal) curves were fit using GraphPad Prism. Means and standard deviations are displayed in the curve fit graphs and were also calculated using GraphPad Prism 8.0.

### Recombinant protein ELISAs

6x-His tag antibodies were coated at 2 ug/mL in PBS onto 96-well half-area high binding plates (Corning, 3690) overnight at 4°C or 2 h at 37°C. After washing, plates were blocked with 3% BSA for 1 h at RT. Then 1 ug/mL of his tagged recombinant SARS-CoV-2 (or SARS-CoV-1) RBD or S proteins were added in plates and incubated for 1 h at RT. After washing, serially diluted antibodies were added in plates and incubated for 1 h at RT. After washing, alkaline phosphatase-conjugated goat anti-human IgG Fcγ secondary antibody (Jackson ImmunoResearch, 109-055-008) was added in 1:1000 dilution and incubated for 1 h at RT. After final wash, phosphatase substrate (Sigma-Aldrich, S0942-200TAB) was added into each well. Absorption was measured at 405 nm.

### Polyspecificity reagent (PSR) ELISAs

Solubilized CHO cell membrane protein (SMP), human insulin (Sigma-Aldrich, I2643), single strand DNA (Sigma-Aldrich, D8899) were coated onto 96-well half-area high-binding ELISA plates (Corning, 3690) at 5 ug/mL in PBS overnight at 4°C. After washing, plates were blocked with PBS/3% BSA for 1 h at RT. Antibody samples were diluted at 100 ug/mL in 1% BSA with 5-fold serial dilution. Serially diluted samples were then added in plates and incubated for 1 h at RT. After washing, alkaline phosphatase-conjugated goat anti-human IgG Fcy secondary antibody (Jackson ImmunoResearch, 109-055-008) was added in 1:1000 dilution and incubated for 1h at RT. After final wash, phosphatase substrate (Sigma-Aldrich, S0942-200TAB) was added into each well. Absorption was measured at 405 nm.

### HEp2 epithelial cell polyreactive assay

Reactivity to human epithelial type 2 (HEp2) cells was determined by indirect immunofluorescence on HEp2 slides (Hemagen, 902360) according to manufacturer’s instructions. In brief, monoclonal antibody was diluted at 100 ug/mL in PBS and then incubated onto immobilized HEp2 slides for 30 min at RT. After washing, one drop of FITC-conjugated goat anti-human IgG was added onto each well and incubated in the dark for 30 min at RT. After washing, cover slide was added to HEp2 cells with glycerol and the slide was photographed on a Nikon fluorescence microscope to detect GFP. All panels were shown at magnification 40x.

### Surface plasmon resonance methods

SPR measurements were carried out on a Biacore 8K instrument at 25°C. All experiments were carried out with a flow rate of 30 μL/min in a mobile phase of HBS-EP [0.01 M HEPES (pH 7.4), 0.15 M NaCl, 3 mM EDTA, 0.0005% (v/v) Surfactant P20]. Anti-Human IgG (Fc) antibody (Cytiva) was immobilized to a density ~7000-10000 RU via standard NHS/EDC coupling to a Series S CM-5 (Cytiva) sensor chip. A reference surface was generated through the same method.

For conventional kinetic/dose-response, listed antibodies were captured to 50-100 RU via Fc-capture on the active flow cell prior to analyte injection. A concentration series of SARS-CoV-2 RBD was injected across the antibody and control surface for 2 min, followed by a 5 min dissociation phase using a multi-cycle method. Regeneration of the surface in between injections of SARS-CoV-2 RBD was achieved by a single, 120s injection of 3M MgCl2. Kinetic analysis of each reference subtracted injection series was performed using the BIAEvaluation software (Cytiva). All sensorgram series were fit to a 1:1 (Langmuir) binding model of interaction.

### Expression and purification of Fab

The CC12.3 Fab was expressed and purified using a previous protocol (Yuan et al., 2020b). In brief, the heavy and light chains were cloned into phCMV3. The plasmids were transiently co-transfected into ExpiCHO cells at a ratio of 2:1 (HC:LC) using ExpiFectamine™ CHO Reagent (Thermo Fisher Scientific) according to the manufacturer’s instructions. The supernatant was collected at 10 days post-transfection. The Fabs were purified with a CaptureSelect™ CH1-XL Affinity Matrix (Thermo Fisher Scientific) followed by size exclusion chromatography. The eCR3022.20 Fab was purified by digesting eCR3022.20 IgG using Fab digestion kit (ThermoFisher, 44985) according to manufacturer’s instructions. After digestion, Fc fragments and undigested IgG were removed from binding to the protein A beads. The unbound flowthrough Fab was collected and followed by size exclusion chromatography.

### Crystal structure determination of the eCR3022.20-RBD-CC12.3 complex

Purified eCR3022.20 Fab, CC12.3 Fab, and SARS-CoV-2 RBD were mixed at an equimolar ratio and incubated overnight at 4°C. The complex (12 mg/ml) was screened for crystallization using the 384 conditions of the JCSG Core Suite (Qiagen) on our custom-designed robotic CrystalMation system (Rigaku) at Scripps Research by the vapor diffusion method in sitting drops containing 0.1 μl of protein and 0.1 μl of reservoir solution. Optimized crystals were then grown in 0.1 M sodium citrate - citric acid buffer pH 5.0, 15% (v/v) ethylene glycol, 1 M lithium chloride, and 10% (w/v) polyethylene glycol 6000 at 20°C. Crystals were grown for 7 days and then flash cooled in liquid nitrogen. Diffraction data were collected at cryogenic temperature (100 K) at the Stanford Synchrotron Radiation Lightsource (SSRL) on the Scripps/Stanford beamline 12-1 with a wavelength of 0.97946 Å, and processed with HKL2000 (Otwinowski and Minor, 1997). Structures were solved by molecular replacement using PHASER (McCoy et al., 2007) with PDB 6XC7(Yuan et al., 2020b). Iterative model building and refinement were carried out in COOT (Emsley et al., 2010) and PHENIX (Adams et al., 2010), respectively.

### Data availability

Crystal structure data and coordinates will be deposited in the PDB prior to publication.

### Animal study

Groups of twelve 6-8 week old Syrian hamsters were put into 6 treatment groups who each received an intraperitoneal (i.p.) infusion of either 10 mg, 2 mg, 0.5 mg, or 0.125 mg per animal of the eCR3022.7 monoclonal antibody or 10 mg per animal of the parental CR3022 monoclonal antibody or 10 mg per animal of an anti-dengue isotype matched control antibody (Den3). After 72 hours, serum was obtained to quantify mAb titers prior to animal infection. Each hamster was then infected through intranasal administration of 10^5 total PFU (plaque forming units) of SARS-CoV-2 (USA-WA1/2020). Animal weights were obtained during the study as a measure of disease progression. On day four post infection, six of the animals were sacrificed and lung tissue was harvested for viral titer analysis by RT-qPCR as well as live viral titers via plaque assay. At day seven post-infection, six of the animals were sacrificed and serum was collected to assess mAb titer at the time of sacrifice using our recombinant protein ELISA protocol. Research protocol was approved and performed in accordance with Scripps Research IACUC Protocol #20-0003.

### Viral load measurements - Plaque Assay

SARS-CoV-2 titers were measured by homogenizing lung tissue in DMEM 2% FCS using 100 μm cell strainers (Myriad, 2825-8367). Homogenized organs were titrated 1:10 over 6 steps and layered over Vero-E6 cells. After 1 h of incubation at 37°C, a 1% methylcellulose in DMEM overlay was added, and the cells were incubated for 3 days at 37°C. Cells were fixed with 4% PFA and plaques were counted by crystal violet staining.

### Statistical methods

Statistical analysis was performed using Graph Pad Prism 8 for Mac, Graph Pad Software, San Diego, California, USA. Groups of data were compared using several methods including the grouped parametric One-Way ANOVA test and the grouped non-parametric Kruskall-Walli test. Data were considered statistically significant at p < 0.05.

